# PHIROS: Integrated microfluidic platform for multi-day high-resolution imaging of organotypic slices

**DOI:** 10.1101/2025.02.28.640791

**Authors:** Jana B. Petr, Meng-Syuan Lin, Catarina M. Gomes, Dina Hochuli, Andreas Hierlemann, Martin Baumgartner, Mario M. Modena

## Abstract

Replicating the cytoarchitecture and cell heterogeneity of the brain *in vitro* remains challenging. Although *ex vivo* organotypic slices preserve native tissue complexity, current culturing methods limit long-term, high-resolution imaging and restrict temporally controlled perturbations. Here, we present PHIROS, a microfluidic platform for high-resolution imaging of organotypic slices that supports static culturing of excised tissue at the air–liquid interface and provides continuous optical access for extended imaging at subcellular resolution. Controlled perfusion with oxygenated medium preserves tissue viability over several days, enabling, e.g., the monitoring of spontaneous and pharmacologically modulated astrocytic calcium activity. Using PHIROS we characterized medulloblastoma (MB) cell behavior in a physiological tumor microenvironment and observed dynamic F-actin-driven interactions with tissue-resident astrocytes, as well as leading edge localization of the immune-checkpoint marker B7-H3 in invading tumor cells. Quantitative assessment of mitochondria transfer across heterotypic actin-rich connections evidences the potential of PHIROS as a versatile system for mechanistic studies in a tissue context, enabling controlled compound exposure and high-resolution imaging in physiologically relevant tissue settings.

## Main

Replicating the complexity of intact tissue *in vitro* remains a major challenge in biology. Intact tissues integrate diverse cell types within a highly organized cytoarchitecture, enabling complex cell signaling and physiological functions. Conventional *in vitro* systems, such as 2D co-cultures, spheroids and organoids, provide valuable insights into cell-to-cell interactions and cellular mechanisms, but fall short of reproducing complex tissue environments, ultimately resulting in limited physiological relevance^1,2^. Consequently, multicellular processes, such as neural network activity, tumor-host interactions, and drug responses, cannot be investigated under conditions that reflect native tissue organization by using these models.

Organotypic tissue slices offer a physiologically relevant alternative to overcome these limitations and preserve cytoarchitecture, cell heterogeneity, cytokine signaling and functional connectivity *ex vivo*^3,4^. These features enable studies of cell motility and signaling within a native tissue context, e.g., in the brain, where neuronal and glial interactions have shown to remain intact^3,5–7^. Organotypic slice models are commonly used in neuroscience and have recently increasingly been applied to cancer neuroscience research^8–12^. The preservation of a neural tumor microenvironment (TME) enables to investigate the interplay between tumor and cells of the TME, where connectivity and heterogeneity strongly influence tumor initiation^13^, progression^14–17^ and metastasis^18^. By engrafting tumor cells on healthy organotypic slices, tumor-host tissue interactions can be studied in a controlled and accessible environment over multiple days^9,19^. While these models can overcome many limitations of conventional *in vitro* systems, the currently used culturing methods introduce restrictions on imaging and experimental flexibility.

Current slice culturing techniques rely on air-liquid interfaces (ALIs)^5^ or alternating exposure to medium and air^20^, which limit continuous optical access to the slice and confine imaging to short time windows or endpoint analysis. Orthotopic tumor models offer restricted imaging and typically require adult animals, large numbers of implanted cells, and long preparation times, constraints that hinder approaches of personalized medicine and raise ethical concerns^10,11^. In contrast, slice-based engraftment requires fewer cells, leverages tissue from young mice that better mimic pediatric tumor biology, and enables short experimental timelines. These advantages make organotypic slices particularly suited for modelling brain tumors, including pediatric brain tumors. such as medulloblastoma (MB), the most common malignant brain tumor in children^21^.

To address slice-culturing limitations and unlock the full potential of slice-based disease modeling, we developed a microfluidics-based **P**latform for **Hi**gh-**R**esolution imaging of **O**rganotypic **S**lices (PHIROS), which features long-term tissue viability, along with continuous, unhindered optical access, and enables precise temporal control of compound delivery. PHIROS enables real-time monitoring of subcellular events, including calcium dynamics, mitochondrial transfer, and actin remodeling over multiple days in a tissue context at unprecedented resolution. To demonstrate potential uses of PHIROS in neuroscience and cancer research, we showcase here two different applications: First, we monitored the calcium (Ca^2+^) dynamics of astrocytic populations in organotypic cerebellar slices over 3 days, and we perturbed network activity by blocking gap-junction intercellular communication. Secondly, we implanted different fluorescently labelled medulloblastoma cells into cerebellar slices and monitored tissue invasion, tumor-tumor and tumor-TME interactions, and the dynamic localization of B7-homologoues 3 (B7-H3), a clinically relevant checkpoint inhibitor, during tissue invasion.

## Results

### Platform for long-term high-resolution imaging of organotypic brain slices

PHIROS integrates four key functions: (i) static culturing at the air-liquid interface (ALI) for tissue recovery and reorganization after slicing, (ii) perfused culturing in a closed chamber for continuous, long-term high-resolution imaging under aseptic conditions, (iii) controlled change of medium conditions for, e.g., drug delivery, and (iv) tissue fixation and recovery for downstream analysis. Once placed on the platform, the brain slices remain undisturbed during culturing and imaging, which prevents mechanical damage through transfers.

The platform comprises four main components (**Supplementary Figure 1A-B)**: i) a polymethylmethacrylate (PMMA)-based slice holder, featuring a microfluidic channel for the perfusion culturing; ii) a hydrophilic 0.4-µm-pore-diameter polytetrafluoroethylene (PTFE) membrane bonded to the slice holder; iii) a PMMA-based support part; iv) an adhesive foil for sealing the assembly and enabling perfusion. The assembly of i) and ii) is further referred to as the “culture chip”. For static culturing, the culture chip is placed on a standard tissue-culture insert, so that medium exchange can be performed following standard culturing procedures by simply transferring the inserts into a well with fresh medium. The chip can then be removed from the insert and sealed before transfer to a microscope stage at the beginning of the perfusion protocol to enable long-term, high-resolution imaging (**Figure 1A-B**).

**Figure 1:**
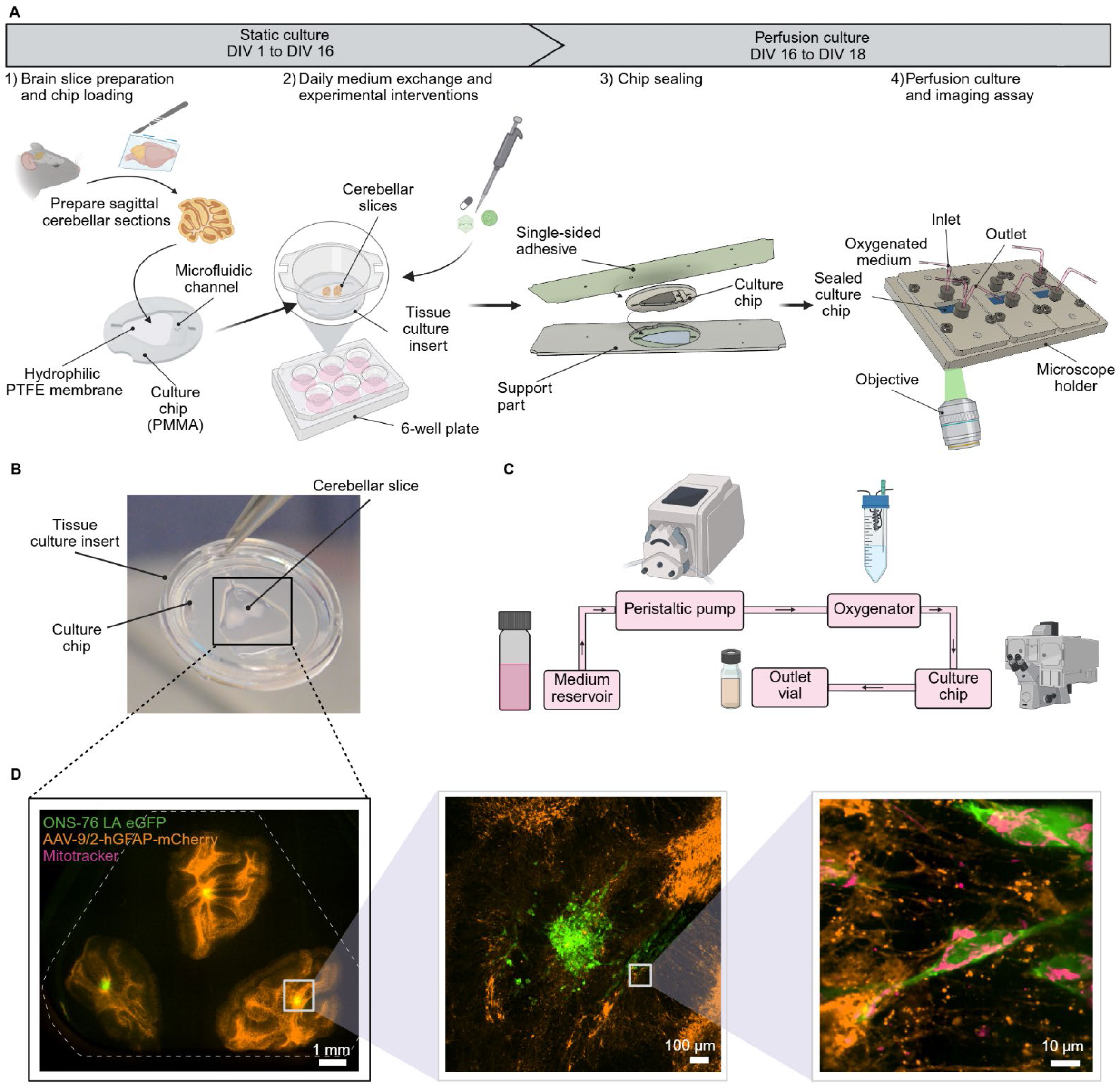
Platform design for long-term high-resolution imaging of organotypic slices. **(A)** Experimental set-up of the platform and workflow using mouse-derived cerebellar slices to investigate cellular dynamics, such as brain tumor motility or intracellular calcium dynamics. An exploded view of the platform design for both static and perfused configuration can be seen in **Supplementary 1.** (**B)** Photo of the developed platform placed on top of a tissue culture insert with two cerebellar slices. **(C)** Schematic of the developed fluidic system to maintain organotypic-slice health on a spinning-disc confocal microscope. A peristaltic pump was used to drive the medium from the main reservoir to the oxygenator (gas-exchanger), buffering the medium to a pH of 7.4, and oxygenating the medium before it reaches the culture chip in the environmental box of the microscope. A detailed view of the oxygenator and the measured dissolved oxygen concentrations can be seen in **Supplementary 2**. **(D)** The developed platform enables to study events across spatial and temporal scales, ranging from 4X or 10X magnification to single-cell dynamics and cell-cell interactions at 60X magnification over multiple days. Images show maximum intensity projections at the respective magnification.

We cultured murine cerebellar slices for 14-18 days at the ALI to enable tissue recovery and Purkinje cell-layer reorganization. ALI incubation enabled experimental interventions, such as adeno-associated-virus (AAV)-based labeling of specific cell subpopulations and medulloblastoma spheroid engrafting onto the cerebellar slices. Imaging was initiated after sealing the slice compartment under sterile conditions (**Supplementary Figure 1B**). Up to 3 chips, accommodating a total of up to 9 cerebellar slices, were perfused and imaged in parallel on a spinning-disk confocal microscope (**Supplementary Figure 1C**). To support brain-slice viability, the medium was oxygenated using a custom-made gas exchanger. We found a dissolved oxygen concentration of ∼30 mg/L being sufficient to support tissue health without bubble formation in the tubing. Oxygenation was achieved by using the custom-made gas exchanger between medium reservoir and chip (**Figure 1C, Supplementary Figure 2**). By combining static and perfusion culturing, PHIROS enables continuous, multi-day monitoring of tissue and single-cell dynamics, including subcellular events, such as actin remodeling and mitochondrial transfer (**Figure 1D**).

### PHIROS preserves cerebellar slice cytoarchitecture and sustains tissue viability

We characterized cerebellar-slice cultures on chip and compared tissue organization with those of standard ALI cultures and *in vivo* metrics, and we assessed tissue health under perfusion. Our main questions were: i) does chip geometry and dual-membrane ALI culture support tissue reorganization, and ii) is viability under perfusion with oxygenated medium maintained.

We cultured P10-P12 murine cerebellar slices either on standard tissue-culture inserts or on the chip placed on an insert and assessed Purkinje cell layer reorganization^8^ after 16 days *in vitro* (DIV 16, **Figure 2A**, dashed line at DIV 16). Under both conditions, slices appeared translucent, indicative of good tissue health and maturation^3^ (**Supplementary Figure 3**). Immunofluorescence staining revealed a continuous Purkinje cell layer along the tissue convolutions, a dense granule cell population, and astrocyte networks in the slices (**Figure 2B, Extended Data 1**). The Purkinje cell layer was flanked by organized granular and molecular layers, indicating a physiologically relevant reorganization of the tissue slice during the static culturing.

**Figure 2:**
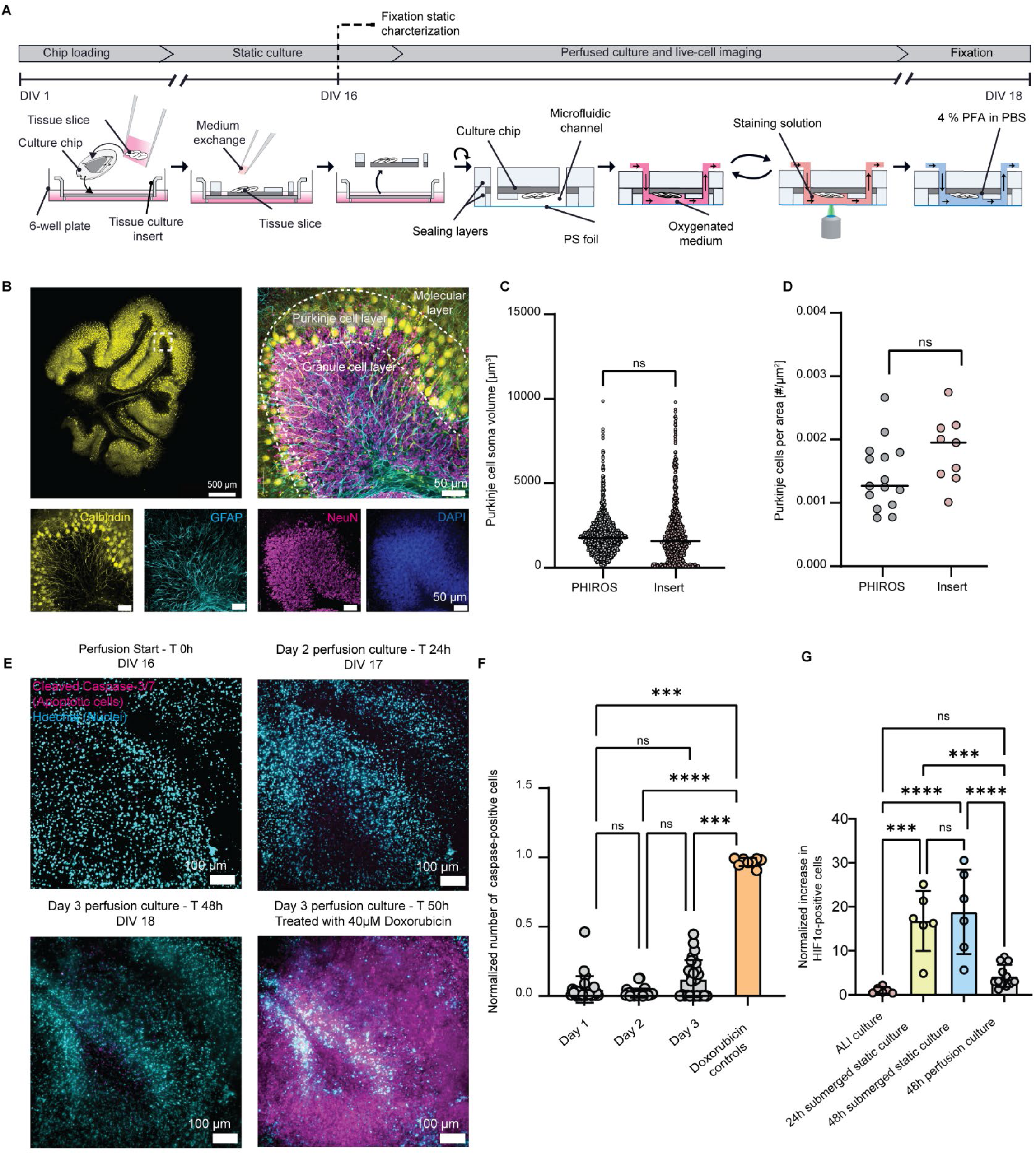
Characterisation of chip-based cerebellar slice cultures and perfusion strategy for long-term imaging. **(A**) Experimental design to assess tissue reorganization during double-membrane culturing (dotted line endpoint at DIV 16) and to determine tissue viability under perfusion at DIV 18. **(B)** Stitched, maximum-intensity projection image of a representative cerebellar slice. The integrity of the Purkinje cell layer marked in yellow using anti-Calbindin is visualized. Composite image of the close-up in **B** with split channels showing markers for anti-Calbindin (yellow), anti-GFAP (cyan), anti-NeuN (magenta) and DAPI (blue). **(C)** Quantitative comparison of Purkinje cell volumes between cerebellar slices cultured on culture chips (n= 6 chips/9 slices, 1658 cells) and on inserts (n=3 inserts, 754 cells). **(D)** Quantitative comparison of Purkinje cell density per area between cerebellar slices cultured on chips (n= 6 chips) and on inserts (n= 3 inserts). Statistics for C and D were derived using an unpaired t-test with Welch’s correction. **(E)** Composite, maximum-intensity projection images of the same slice stained for nuclei (cyan) and apoptotic cells (cleaved caspase 3-7, magenta) after perfusion start, 24h into the perfusion culture, 48h into the perfusion culture, and 2h post exposure to 40 µM doxorubicin. **(F)** Quantitative assessment of caspase-positive cells (n = 6 chips/12 slices). **(G)** Quantification of hypoxic cells by using immunostaining for anti-HIF1-alpha and nuclear co-localization. Results are normalized with respect to hypoxic-cell counts found in standard ALI cultures. N = 4 slices per condition. Statistics for F and G were derived using 1-way ANOVA.

We next quantified Purkinje cell-soma size and cell density and compared these values to those of standard ALI cultures and published *in vivo* data. Early culture chip prototypes with suboptimal culture conditions yielded sparse, disorganized, and disrupted Purkinje cell layers with enlarged cell somas (**Supplementary Figure 4**). In contrast, the Calbindin staining of the final chip design revealed a mean Purkinje cell-soma area of 2028 µm^3^ and a cell density of 0.0014 cells/µm^2^, matching data of standard ALI cultures and previously reported values for Purkinje cell density *in vivo*^22^ (**Figure 2C-D**). These results confirmed that chip geometry and dual-membrane culture supported the reorganization of the cerebellar architecture in organotypic slices without compromising viability.

To perform high-resolution imaging, the tissue slices need to be submerged in medium. However, submerged slices develop hypoxia in the absence of perfusion, and imaging is restricted to short time periods of a few hours only. To overcome this limitation, we implemented a pulsatile-perfusion protocol with oxygenated medium, designed to minimize disruption of endogenous gradients in the culture chamber while ensuring sufficient oxygen supply. We established a flow protocol consisting of hourly cycles of partial medium exchange with intermittent 1 h no-flow intervals. This strategy resulted in a daily medium consumption of ∼1.4 mL, a volume comparable to that used under standard culturing conditions (1 mL) (**Figure 2A**).

To assess cell viability under perfusion, we used a cleaved caspase-3/7 live-cell stain, which visualizes apoptotic cells in real time. Cell viability was assessed at the start of the perfusion (day 1), after 24 (day 2) and 48 h (day 3). As a positive control, slices were treated with 40 µM doxorubicin after 48 h of perfusion to induce apoptosis (**Figure 2E, Extended Data 2A**). The low number of caspase-3/7-positive cells prior to doxorubicin exposure indicated good tissue health in the sealed chip throughout the 48 h of perfusion (**Figure 2F)**. No significant increase in apoptotic cells was observed during the perfusion period, which further confirmed that medium oxygenation and pulsatile perfusion supported cell viability.

To evaluate hypoxic stress during perfusion culturing, we determined the expression levels of the hypoxia transcription factor HIF1-α in the tissue slices by immunofluorescence analysis. We compared the cerebellar slices cultured using the pulsatile flow protocol with slices either cultured at the ALI or submerged for 24 or 48 hours under 2 ml of medium (∼2 mm of medium column above the slices), to mimic the conditions of continuous, long-term imaging using immersion objectives. Nuclear localization of HIF1-α, indicative of hypoxia, was quantified and normalized against control ALI conditions. Submerged slices under static conditions exhibited a 16.8- and 18.8-fold increase in hypoxia levels after 24 and 48 hours, respectively. In contrast, hypoxia levels under pulsatile perfusion with oxygenated medium did not show any significant difference with respect to ALI control condition (**Figure 2G, Extended Data 2B**). These findings indicate that slice submersion, which would be required for imaging using high numerical-aperture objectives, would rapidly compromise slice conditions, while pulsatile flow with oxygenated medium in a closed environment preserves slice viability, thereby enabling long-term imaging investigations.

### Multi-day imaging of subcellular and network-wide astrocytic calcium events

To demonstrate PHIROS’ ability to perform functional imaging in an intact tissue context, we monitored spontaneous astrocytic Ca^2+^ dynamics in cerebellar slices. Astrocytes form a network that is highly interconnected through gap junctions, enabling the transfer of ions and small molecules between cells. In response to neuronal excitation, astrocytes exhibit complex intracellular Ca^2+^ fluctuation and global Ca^2+^ dynamics. These calcium changes can be visualized using genetically encoded calcium indicators^23,24^ and detected using fluorescence microscopy.

We transduced cerebellar slices from P8-P10 mice with AAV-hGFAP-GCaMP6f and maintained the cultures at the ALI for 10 days (**Figure 3A)**. During calcium imaging, we transiently replaced the culture medium with artificial cerebrospinal fluid (aCSF) using the flow system. To support tissue viability, standard culture-medium conditions were reinstalled between imaging sessions. PHIROS enabled repeated imaging of the same region of interest (ROI) over three consecutive days, allowing long-term monitoring of astrocytic activity in a physiologically relevant tissue context (**Figure 3B-D**, **Supplementary Video 1**).

**Figure 3:**
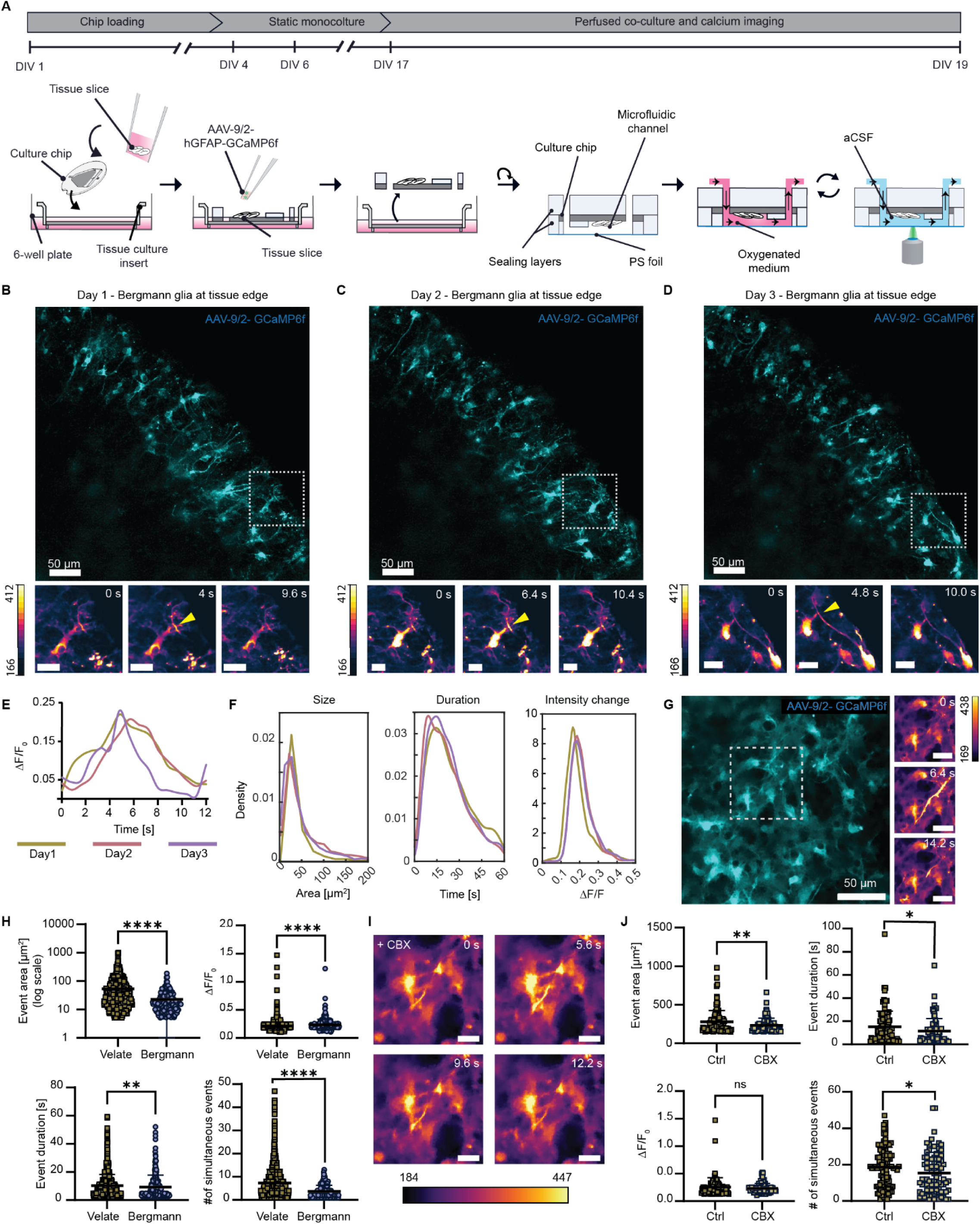
Multi-day monitoring of spontaneous calcium dynamics in cerebellar tissue. **(A)** Experimental protocol to investigate intracellular calcium changes in cerebellar astrocytes; the static culturing period was used to label astrocytes via AAV transduction. **(B-D)** Images of the same ROI during the calcium imaging assay on day 1, day 2 and day 3 of the perfusion culture. Close-ups show a calcium event indicated by the yellow arrow in the astrocytic process of likely the same cell across the three time points. Scale bar of the close-ups is 20 µm. **(E)** Fluorescence intensity changes of the events depicted in **B-D**. **(F)** Detected changes in calcium event area, duration, and changes in fluorescence intensity from day 1 to day 3 of the imaging assay (n = 4 slices from 2 independent experiments, 3-4 ROIs per slice), visualized as probability density functions. **(G)** Representative example of calcium dynamics of velate astrocytes. The insets show a calcium event likely traveling across multiple cells in the astrocyte network. **(H)** Comparison of different calcium-event metrics between velate and Bergmann glia cells (n= 4 slices/ 2 chips each). **(I)** Calcium event in the same ROI as in G after carbenoxolone exposure. **(J)** Comparison of different metrics of calcium events in velate astrocytes before and after carbenoxolone exposure.

We detected spontaneous intracellular Ca^2+^ signal fluctuations with consistent intensity changes (∼0.2 ΔF/F) in astrocytic processes across the three days (**Figure 3E**). Ca^2+^ event characteristics were analyzed using AQuA2^25^, including event duration, area, and intensity, and remained stable throughout the 48 hours of perfusion culture (**Figure 3F**), confirming the platform’s ability to support tissue functionality and facilitate functional imaging over extended periods.

Next, we investigated the difference in Ca^2+^ events between Bergmann glia (**Figure 3D**) and velate astrocytes (**Figure 3G**), two cerebellar astrocyte subtypes that display distinct morphological and molecular signatures and local connectivity^26^. Bergmann glia cells extend radial processes across the Purkinje cell layer to the pial surface, while velate astrocytes are distributed within the granule cell layer^27^. We observed Ca^2+^ fluctuations in the cell body and in processes in both astrocyte subtypes. Notably, we observed Ca^2+^ waves spreading within local astrocyte networks predominantly in regions rich in velate astrocytes (**Figure 3G, Supplementary Video 2**). While further investigating the Ca^2+^ dynamics of these two astrocyte subtypes, we found that Ca^2+^ events in Bergmann glia were smaller in size, showed a smaller amplitude and were shorter than observed events in velate astrocytes. Furthermore, the number of events detected simultaneously was higher in velate astrocyte regions, indicating higher cellular connectivity (**Figure 3H**). We next assessed whether velate network activity could be modulated acutely using the gap-junction inhibitor carbenoxolone (CBX). Following exposure, spontaneous Ca^2+^ fluctuations persisted, but event areas became smaller, likely including single cells or small clusters of neighboring cells (**Figure 3I**). Quantification revealed a decrease in the size and duration of events covering an area larger than 150 µm^2^, as well as a reduction in simultaneously occurring events, indicating carbenoxolone-induced disruption of gap-junction-mediated network communication. Together, our findings underline PHIROS’ ability to support multiscale Ca^2+^ event visualization at sub-cellular resolution and over multiple days, while enabling precise control over medium conditions at high temporal resolution.

### Actin cytoskeleton dynamics and tissue-invasion behavior of medulloblastoma cells

Tumor cell invasion is strongly influenced by tissue stiffness and composition, features that differ between *in vivo* and classical *in vitro* migration models. Tissue composition forces cells to dynamically adapt motility modes during microenvironmental exploration^28^. We used PHIROS to capture time-lapse recordings of MB spheroids, co-cultured with cerebellar slices, and evaluated TME-dependent changes in MB cell morphology and motility by comparing them to tumours grown on dissociated neural cells. During the static culturing period of the cerebellar tissue slices, we transduced the slices with hGFAP-driven mCherry, enabling the visualization of astrocyte somata and processes. To simultaneously monitor F-actin dynamics, we used ONS-76 cells expressing LifeAct-eGFP. Tumor spheroids were placed on slices 72 h before imaging (DIV 16–19) to allow engraftment, and sealed chips were imaged for 48 h under periodic perfusion (**Figure 4A**).

**Figure 4:**
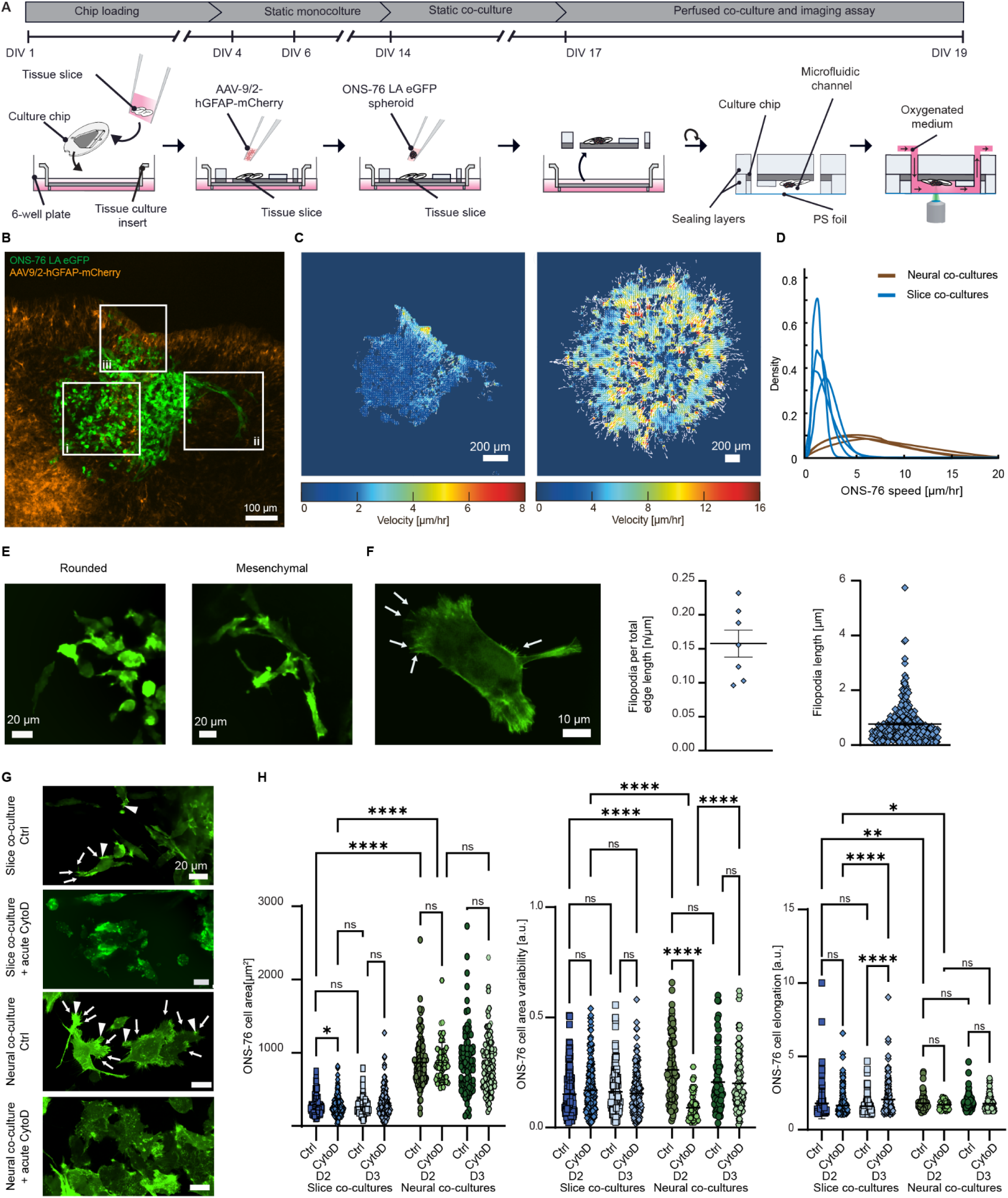
Dynamics of cell flow and single-cell trajectories in ONS76-LAeGFP spheroids. **(A)** Experimental workflow to establish the medulloblastoma co-culture model for high-resolution imaging assays. **(B)** Representative image showing the ONS-76-LA-eGFP cells and astrocytes during 12-hour time-lapse imaging. Green: LA-eGFP; orange: GFAP-driven mCherry. Areas *i* and *iii* exhibit increased cell motility, while area *ii* shows confined cellular movement. **(C)** Heat maps display the mean velocity magnitude across the spheroid invading a slice (left) or migrating on dissociated neural cells (right) over 12 hours. Arrows indicate the mean motion direction. **(D)** Fitted mean velocity magnitudes from **(C)** plotted as a density distribution (n = 4 spheroids from slice co-cultures and n=3 spheroids from dissociated neural cell co-cultures) **(E)** Representative images of tumor cells invading slices displaying rounded and mesenchymal-like invasion phenotypes **(F)** Representative image of the actin-cytoskeleton of an invading medulloblastoma cell in the cerebellar slice. Arrows point to filopodia extending from the cell body. Quantifications show the normalized number of filopodia and filopodia lengths of MB invading the tissue (n= 12 cells from 4 slices/ 2 chips) **(G)** Representative images of untreated and 1 µM cytochalasin-D treated ONS-76 cells co-cultured with cerebellar tissue slices or neural dissociated cells. Arrowheads indicate exemplary TME-exploring lamellipodia, arrows filopodia in untreated cells. **(H)** ONS-76-LA-eGFP cell area, cell area variability, and cell elongation were assessed at day 2 under standard medium conditions and during CytoD exposure (D2 Ctrl and D2 CytoD, respectively), and 24h after exposure (D3 Ctrl and D3 CytoD). N = 4 control slices, n = 5 CytoD slices, n = 5 ROIs control neural co-cultures, n= 3 ROIs CytoD treated neural co-cultures). Nonparametric one-way ANOVA was used to derive statistical differences.

We found that the tumor cells stopped exploring the cerebellar tissue microenvironment under continuous perfusion, indicating that the flow might interfere with microenvironmental factor gradients. This observation highlights the immediate impact of perturbation and underscores the importance of the microenvironment for invasion. To minimize the disruption of endogenous gradients, we limited the flow to only 7 minutes every hour, while image acquisitions were performed outside the flow intervals. Under these flow conditions, we could observe different patterns of motile cell behavior, ranging from random motility in the spheroid center (**Figure 4B i**) to directional streams of collectively invading cells (**Figure 4B ii**). The invading cells exhibited directional movement trajectories only in certain regions of the tumor cell mass on the tissue slice. In contrast, ONS-76 cells migrated radially on dissociated cultures of neural cells without constraining tissue barriers (**Supplementary Figure 5**). Particle image velocimetry (PIV) showed spatial heterogeneity in tumor cell dynamics when comparing the 2D with the 3D TME (**Figure 4C, Supplementary Figure 6**). The velocity magnitude analysis revealed uniform radial movements in the neural cell-tumor co-culture, whereas corresponding directional movements in the tissue slices were spatially restricted (**Figure 4C**). Speed analysis revealed higher speeds of cells seeded on neural co-culture compared to speeds on slice co-cultures, the latter of which were in good accordance with *in vivo* values^29,30^, suggesting that the microenvironment architecture served as an important factor controlling tumor motility **(Figure 4D)**.

In the slice co-culture models, the largest velocities were recorded in cluster *iii*, where cells displayed actin-rich structures that were associated with altered local velocity patterns of mCherry-positive astrocytes (**Supplementary Figure 6**), suggesting a potential physical interaction of these two cell populations. Cluster *i* cells display marked F-actin dynamics with little discernible directional movement of the mostly rounded cells (**Figure 4E)**. In contrast, cluster *ii* cells displayed flat and elongated morphologies with prominent lamellipodia and filopodia, reminiscent of mesenchymal cells^31,32^ at the invasion front (**Figure 4E, Supplementary Video 4**). These cells exhibited F-actin-rich lamellipodia that displayed explorative contacts with surrounding cells via filopodia-like protrusions (**Figure 4F**). Importantly, while tumor cell-cell contacts in clusters *i we*re mostly short lived (0.5 – 6 h), the tumor cell-cell contacts in cluster *ii* persisted for more than 12 h and were an indication of collective invasive behavior. We also observed fewer contacts between rapidly invading tumor cells of this cluster with mCherry-positive astrocytes, suggesting an avoidance mechanism that allows tumor cells to navigate through glial structures during invasion. (**Extended Data 3**).

To assess whether acute treatment effects could be monitored using PHIROS, we monitored the effects of actin-polymerization inhibition by exposing both the slice and neural co-cultures to cytochalasin D (CytoD) for 1 hour. In both models, subcellular actin distribution changed after exposure, with a marked loss of F-actin and increased punctate actin distribution **(Figure 4G)**. Furthermore, in neural co-cultures, CytoD induced an acute inhibition of cell dynamics, which was not displayed in slice settings. Conversely, CytoD-treated tumor cells on cerebellar slices elongated markedly within 24 h of exposure, possibly resulting from tissue-associated responses to CytoD **(Figure 4H)**.

### B7-H3 localized to leading edge of explorative and invasive tumor cells

We next investigated whether PHIROS could be utilized to study the behavior of tumor-associated surface molecules within a tissue context. To this end, we engineered ONS-76 cells to express the transmembrane protein B7-H3/CD276, fused with mNeon-Green (ONS-76-B7-H3-mNG, **Figure 5A**). B7-H3 is highly expressed in malignant pediatric brain tumors, including MB, and is a validated target for CAR-T cell therapies^33,34^. In tissue-invading tumor cells, B7-H3-mNG was prominently localized at the leading edge and along the plasma membrane (**Figure 5A-B**). We observed both dynamic and static cell-cell contacts. Dynamic contacts were defined as cell-cell interactions that formed during an observation period of 30 min, whereas static contacts were those that persisted throughout the 30-min period. A rapid increase in B7-H3 intensity was detected at the contact sites of dynamic cell-cell interactions, whereas the B7-H3-mNG signal at static contacts remained relatively stable (**Figure 5C**). Quantification of relative mNG fluorescence across the lamellipodia of invading cells confirmed a pronounced enrichment of B7-H3 at the leading edge (**Figure 5D**). This polar distribution and accumulation were maintained in individual cells for at least 24 hours (**Figure 5E**).

**Figure 5:**
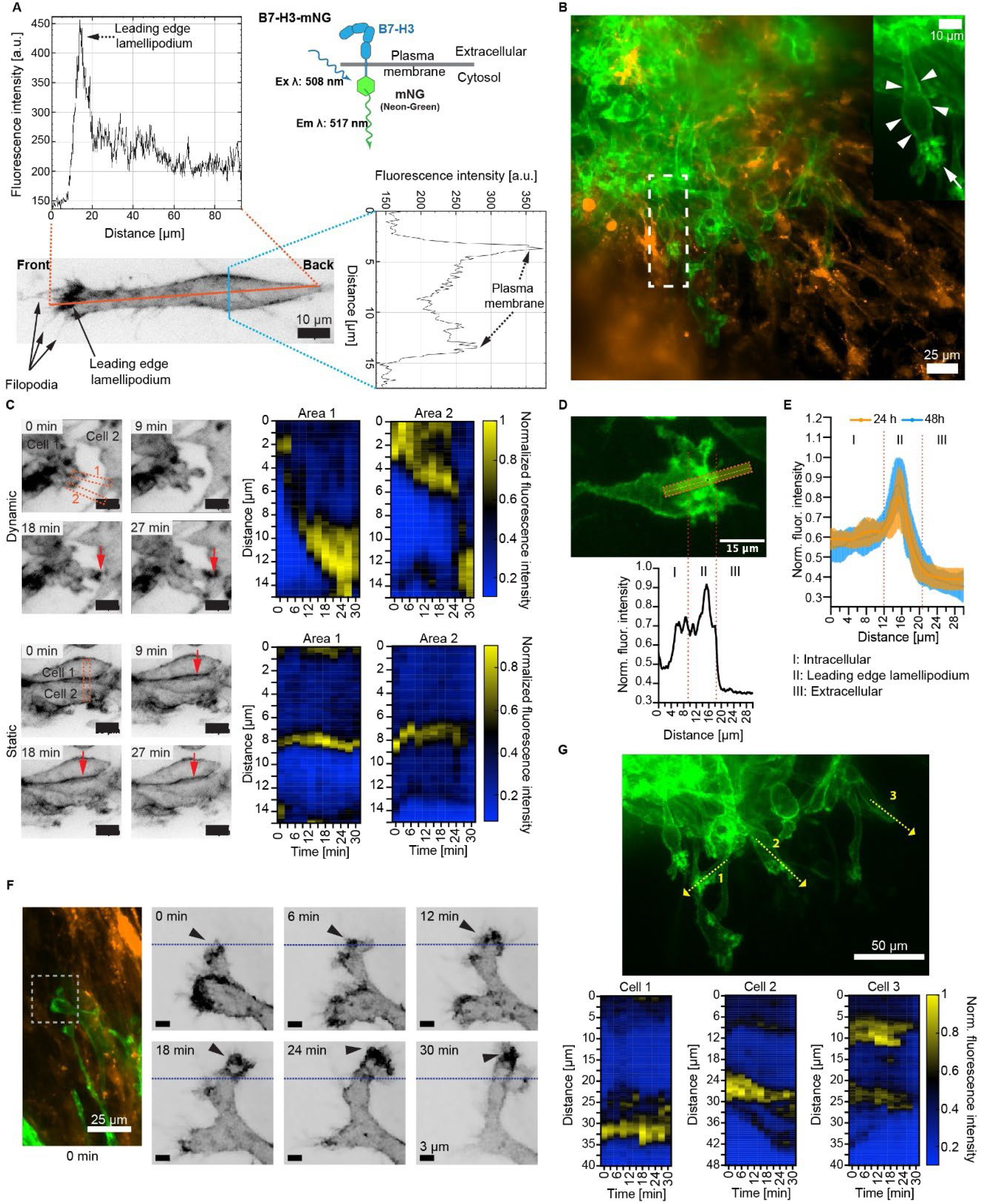
Persistent accumulation of B7-H3 at the leading edge of tissue-invading tumor cells. **(A)** Grayscale image of a polarized ONS-76-B7-H3-mNG cell on a tissue slice during perfusion culturing, two days after chip closure. The plot above the cell shows fluorescence values along the longitudinal profile (orange, horizontal line), while the plot on the right shows values of the cross-sectional profile (blue, vertical line). The schematic at the top right illustrates the B7-H3 fusion construct. **(B)** Maximum intensity projection of a confocal image stack after 48 hours of perfusion culture. Green: B7-H3-mNG; orange: GFAP-driven mCherry. The inset highlights the green channel, showing membrane-localized B7-H3 (arrowheads) and the leading-edge lamellipodium (arrow). **(C)** Dynamics of B7-H3-mNG at dynamic (top) versus static (bottom) cell-cell contacts. Heatmap kymographs show normalized fluorescence values along a 15-µm line (yellow dotted line) at cell-cell contacts, with arrows indicating B7-H3 accumulation. **(D)** Representative B7-H3-mNG fluorescence in tissue-invading tumor cells. Below: Normalized values of fluorescence intensity along a 30-µm line. **(E)** Normalized fluorescence values across leading-edge lamellipodia in cells invading during 24 hours (n=17) and 48 hours (n=21). **(F)** Left: Maximum intensity projection of confocal imaging after 48 hours of perfusion culture (green: B7-H3-mNG, orange: GFAP-driven mCherry). Right: Inverted grayscale of B7-H3-mNG fluorescence intensity shows lamellipodia dynamics over 30 minutes, with arrowheads indicating accumulation at the leading edge, whereas the initial position of the leading edge is visualized with a blue dotted line **(G)** Top: Green channel of image in B, with yellow-dotted arrows marking quantified areas of leading-edge lamellipodia. Bottom: Heat map kymographs show relative fluorescence values over 30 minutes along a 40–48 µm line across leading-edge lamellipodia of representative invading cells.

The persistent accumulation of B7-H3 at the leading edge of invading cells and at the tip of TME-exploring lamellipodia indicates a critical role of B7-H3 in lamellipodial guidance and tumor invasion (**Figure 5G, Extended Data 4**).

### Long-term, high-resolution imaging with PHIROS to monitor cell-cell interactions and organelle trafficking in tissue slices

With the established methodology, we were able to repeatedly image the same MB-astrocyte clusters over three consecutive days without any loss of registration or resolution (**Figure 6A**). This stability enabled morphological quantification of astrocyte features, including branching parameters and astrocyte area (**Figure 6B-C**). Our analysis revealed an initial decrease in astrocyte area, followed by an increase during which astrocytes reached similar area values as at the assessment on the first day of the imaging assay. Notably, the number of branches decreased over the three-day observation period, while the branch lengths increased. These morphological changes could indicate an initial, transient state of astrocyte reactivity, triggered by the proximity of the tumor cells, as has been observed in neurological injuries^35^. Furthermore, we observed astrocytic death in proximity to the tumor cells, likely caused by tumor-astrocyte interactions (**Extended Data 5**).The ability to observe these morphological alterations underlines PHIROS’ capability to provide robust structural analysis in a complex tissue environment.

**Figure 6:**
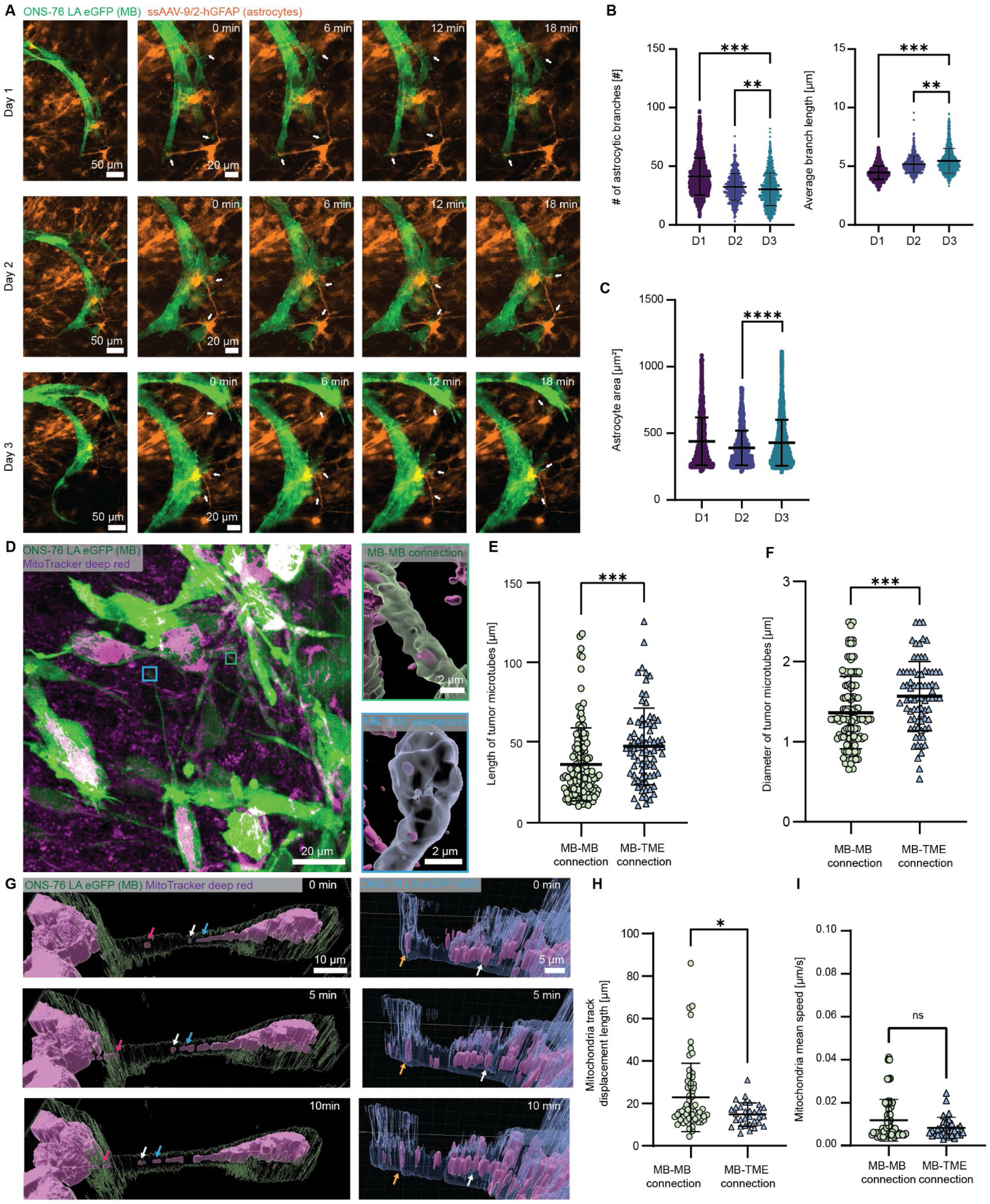
Cell-cell interactions and mitochondria trafficking in a tissue context. **(A):** Interactions of the same cluster of ONS-76-La-eGFP medulloblastoma cells (green: LA-eGFP) with astrocytes (orange: GFAP-driven mCherry) over three days of perfusion culturing and imaging. Images are maximum intensity projections. **(B)** Number of branches and average branch length of astrocytes in close proximity to the tumor. **(C)** Cell areas of astrocytes in close proximity to the tumor cells. Data in (B) and (C) was derived from astrocytes of three different chips (n = 955 cells for D1, 702 cells for D2 and 1334 cells for D3 from 4 slices). Statistical differences between days were derived using a one-way ANOVA. **(D)** Representative maximum intensity projection image of a region featuring actin-rich tumor microtubular connections (green: LA-eGFP) and mitochondria (magenta: MitoTracker deep red). Insets show 3D renderings of a tumor-tumor connection (framed in green) and a tumor-TME connection (framed in blue). **(E)** Length and **(F)** diameter of the tumor-tumor connections (n = 124 TMs from 6 chips/14 slices) and tumor-TME microtubes (connections between medulloblastoma and cells of the TME (blue triangles, n = 78 connections from 6 chips). **(G)** 3D renderings of different timepoints showing a mitochondria (magenta: MitoTracker deep red) transfer between two connected medulloblastoma cells (green: La-eGFP) and in a tumor-TME microtube (blue: LA-eGFP) protrusion connecting to the TME (in blue: LA-eGFP). Colored arrows indicate the movement of the same mitochondria between frames. **(H)** Displacement and **(I)** speed of mitochondria in tumor-tumor connections (n=61 from 6 chips/14 slices) and tumor-TME connections (n=31 from 6 chips/14 slices. Data in G, H, J, K, L was first tested for normal distribution before applying either a Welch’s t-test for lognormal distributed data or a Mann-Whitney t-test.

In our acquisitions, we regularly observed microtubular connections between tumor cells and cells of the cerebellar slices (**Figure 6D**). Some of these connections displayed moving mitochondria, indicating mitochondrial transfer activities across F-actin-positive cell-cell connections (**Figure 6G, Extended Data 5**). In pediatric and adult glial tumors, these connections, referred to as tumor microtubes, generate interconnected networks of cells implicated in therapy resistance^36,37^. Recently, they were also identified in patient-derived xenografts of Li-Fraumeni SHH human medulloblastoma^38^. We observed that tumor-tumor connections were shorter and thinner compared to the microtubes linking tumor cells and cells of the microenvironment (**Figure 6E-F**). While the connections remained stable for several hours, we observed a few events where cells disconnected, or tubes extended towards other cells (**Extended Data 5**). We observed mitochondria moving through these microtubes between tumor cells, a phenomenon not yet addressed in MB, with displacements of 22.8±16.1 µm and a mean speed of 0.012±0.01 µm/s (**Figure 6H-I**).

## Discussion

Organotypic slices retain native cytoarchitecture and multicellular interactions for weeks ex vivo, offering an advantageous model for studying complex cellular dynamics that dissociated cultures or organoids cannot faithfully reproduce. Yet the fact that slices require ALI culturing has prevented continuous high-resolution imaging. PHIROS directly addresses this limitation by enabling multiday, subcellular-resolution imaging under controlled perfusion while maintaining tissue viability. The platform integrates ALI culturing for tissue recovery and extends experimental options, including AAV transduction and spheroid engraftment, while pulsatile-perfusion culturing can be performed in a transparent, sealed chamber that is compatible with high-NA objectives. The platform design ensures stable registration, aseptic conditions, and, by relying on inert, biocompatible plastics, it eliminates adsorption issues associated with soft polymers^39^.

Cerebellar slices retained an intact Purkinje cell layer and preserved laminar organization, confirming tissue health. Although validated for the cerebellum, the developed protocols are applicable to other brain regions, such as hippocampus or thalamus, which are challenging to image dynamically *in vivo*^40^.

PHIROS enabled long-term visualization of spontaneous Ca²⁺ activity at subcellular resolution, revealing astrocytic Ca^2+^ heterogenic dynamics. Pulse delivery of a gap junction inhibitor rapidly disrupted network activity, demonstrating the platform’s capacity to probe acute and sustained responses to targeted perturbations under physiological tissue conditions *in vitro*.

We further used PHIROS to dissect MB invasion in a tissue context. Tumor cells on organotypic slices recapitulated patient and *in vivo*–like phenotypes more faithfully than in neuronal co-cultures. The platform uncovered heterogeneous invasion behavior and helped to resolve complex F-actin polymerization dynamics in invading and non-invading cells, illustrating how MB cells sense and respond to the TME. Single-cell analyses further identified behaviorally distinct MB clusters shaped by local cues. Tracking the transmembrane receptor B7-H3 revealed its persistent accumulation in tumor cell lamellipodia during invasion, suggesting an additional role of B7-H3 in TME interaction and motility beyond its known immunomodulatory functions^41^.

Finally, PHIROS revealed dynamic remodeling of the surrounding TME, including astrocytic processes, and enabled real-time visualization of actin-rich microtubes connecting MB cells, which mediates organelle exchange, a process which has been linked to therapy resistance of gliomas. The ability to track mitochondria trafficking demonstrates that PHIROS provides the resolution and stability that are required to capture subcellular organelle dynamics over multiple days, which is pivotal for studying intercellular communication in brain tumors and neural diseases.

Taken together, our results establish PHIROS as a robust and broadly applicable platform for long-term, high-resolution interrogation of cellular dynamics in intact physiological tissues, enabling mechanistic studies that are not feasible with conventional *in vitro* or *in vivo* approaches.

## Acknowledgements

This work was financially supported by Innosuisse – the Swiss Innovation Agency-under grant 120.665 IP-LS. The authors acknowledge the Single Cell Facility (SCF) at D-BSSE of ETH Zürich for help and support, as well as Dr. Sreedhar Kumar and Dr. Michele Nava, also at D-BSSE, and Dr. Bernard Ciraulo, University Children’s Hospital Zürich, for support with data interpretation and visualization.

## Author contributions

J.B.P. designed and fabricated the platform, designed and performed all experiments, analyzed and interpreted tissue reorganization, tissue viability, hypoxia, calcium imaging, tumor microtube and mitochondria trafficking data, pre-processed the tumor invasion data, and wrote the manuscript. M.S.L designed and performed experiments, interpreted tissue reorganization, analyzed tumor invasion data, and edited the manuscript. C.M.G. analyzed and interpreted astrocyte morphology. D.H. generated and validated the fluorescent cell lines. M.B. conceived the project, acquired funding, provided experimental advice and supervision of the experiments, analyzed tumor invasion and B7H3 accumulation data, and edited the manuscript. A.H. supervised the project, provided experimental advice, and edited the manuscript. M.M.M. conceptualized the project, acquired funding, supervised the project, designed the platform, supported with experimental design and experiments, analyzed tissue viability and tumor invasion data, and edited the manuscript. All authors read and approved the final version of the manuscript.

## Extended Data

**Extended Data 1:**
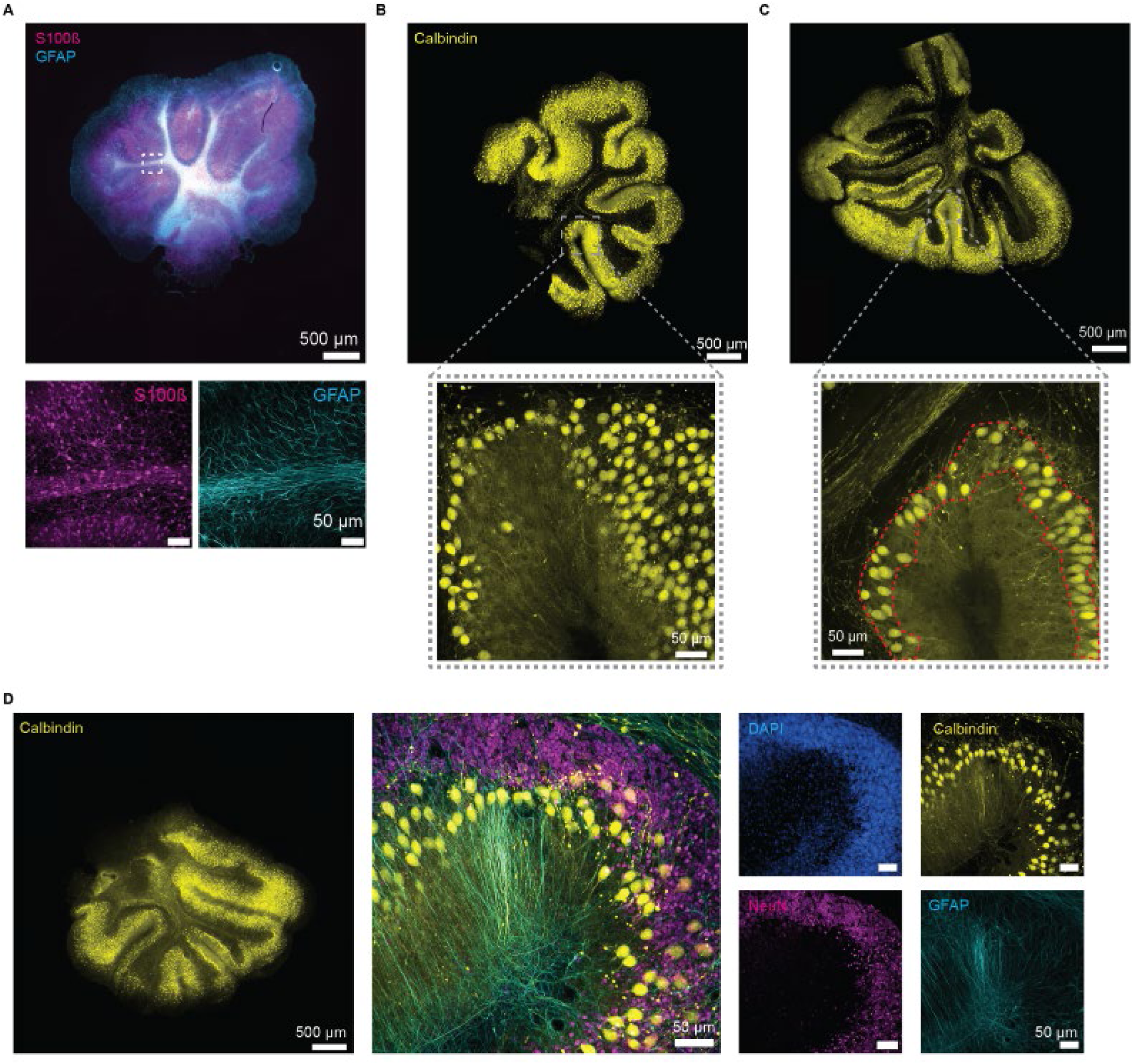
**(A)** 10X maximum intensity projections of the astrocytic networks labeled with anti-S100β in magenta and anti-GFAP in cyan. Close-ups show 40X maximum intensity projections of the same, representative cerebellar slices cultured on PHIROS. (**B-C**) 10X maximum intensity projections of the Purkinje cell layer of slices cultured on PHIROS and stained with anti-Calbindin after fixation at DIV 14 of two additional experiments. Close-ups show 40X images of the Purkinje cell reorganization. The red line in close-up **C** indicates how the boundaries of the Purkinje cell layer were defined to assess the Purkinje cell density. (**D**) Purkinje cell reorganization after 14 days of culturing during which slices were cultured on standard tissue-culture inserts. The close-ups at 40X show the presence of different neuronal cell types, identified by the anti-Calbindin and NeuN staining, as well as glial cells, as indicated by the anti-GFAP staining.

**Extended Data 2:**
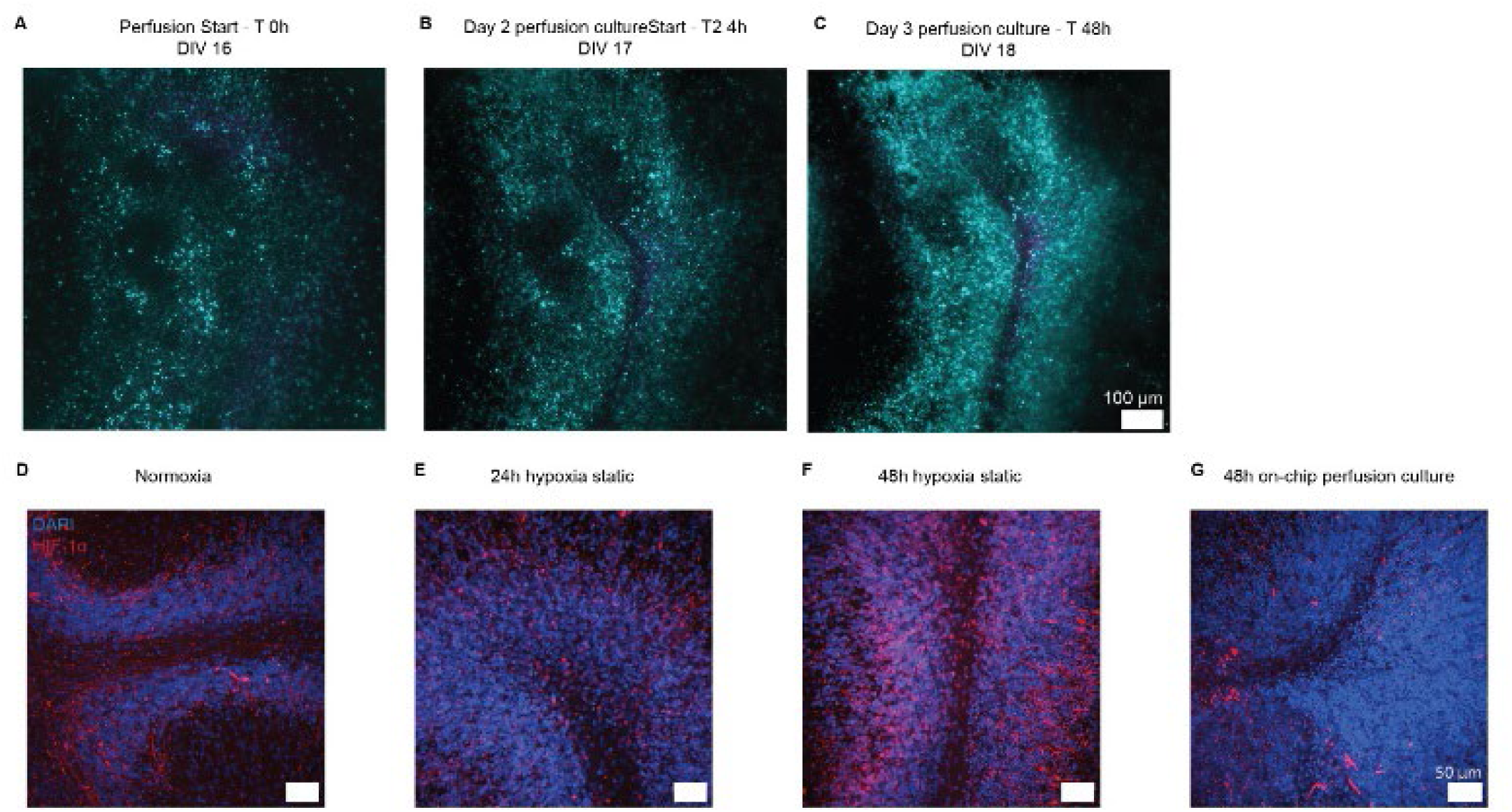
ROIs of different slice experiments of Figure 2. **(A-C)** Viability assessment using cleaved Caspase-3/7 stain labeling apoptotic nuclei (magenta) co-stained with NucBlue (cyan). The images show the same tissue slice during 48 h of perfusion culturing. **(D-F)** Representative immunofluorescent staining for transcription factor HIF1-α (red) and nuclei (DAPI, in blue) of ALI cultured slices (**D**), slices cultured for 24h (**E**) or 48h (**F**) submerged under 2 ml of medium, or for 48h under perfusion on PHIROS (**G**).

**Extended Data 3:**
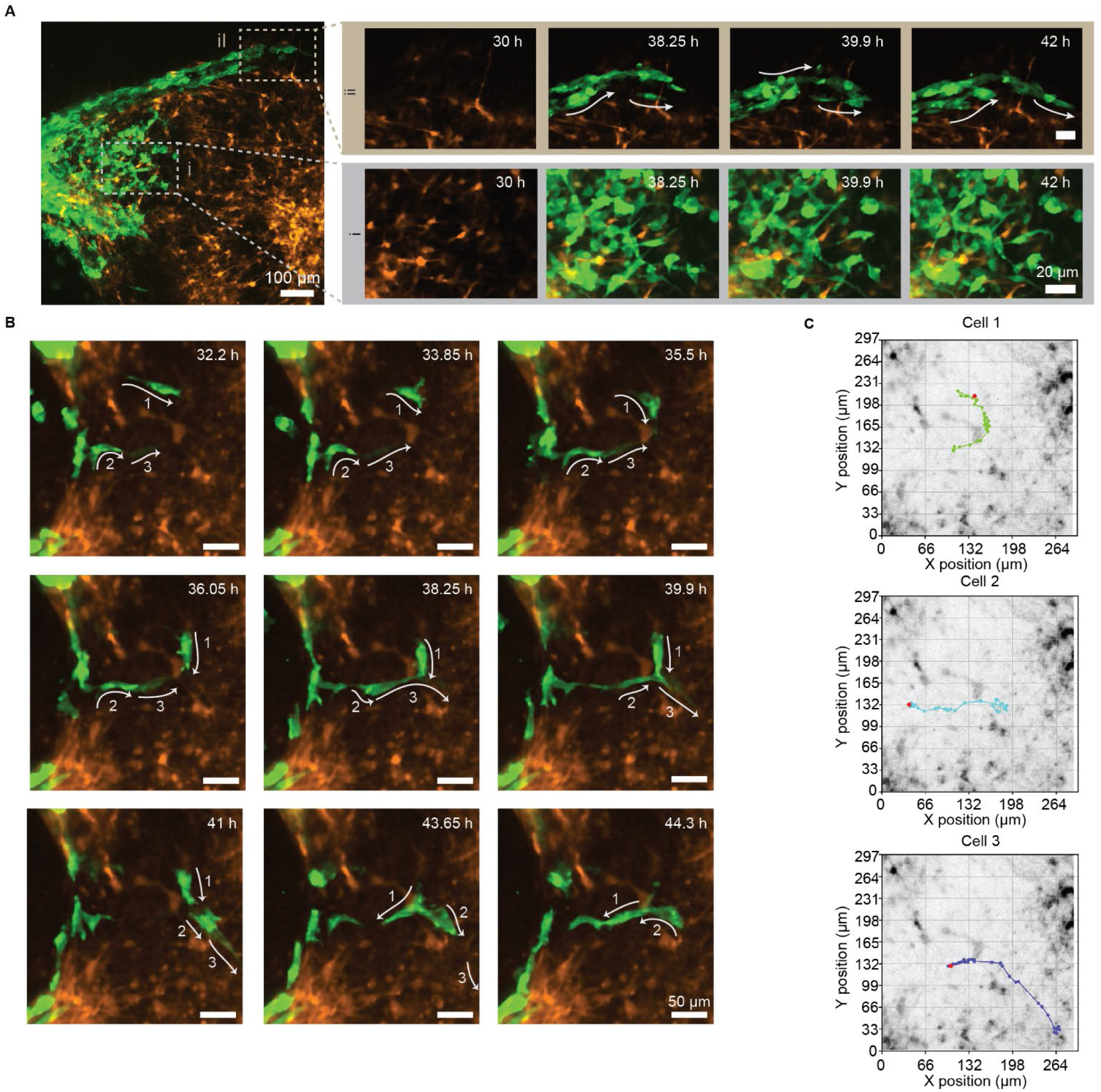
**(A)** Representative image showing tumor-astrocyte interactions during 12-hour time-lapse imaging. Area *ii* highlights a region with low astrocyte density (orange: GFAP-driven mCherry), where pronounced ONS-76-La-eGFP cell movement (green: LA-eGFP) was observed (arrows indicate movement direction). Area *i* shows higher astrocyte density, where cell movement was minimal. Time refers to time after platform sealing; the scale bar of both sets of insets is 20 µm; **(B)** Example of repulsive and attractive interactions between tumor cells (green) and astrocytes (orange) over 14 hours. Movements of cells 1, 2, and 3 are shown with arrows representing their respective trajectories. **(C)** Individual cell trajectories on a background image show only astrocytes at the imaging starting point, illustrating movement patterns in relation to astrocyte density.

**Extended Data 4:**
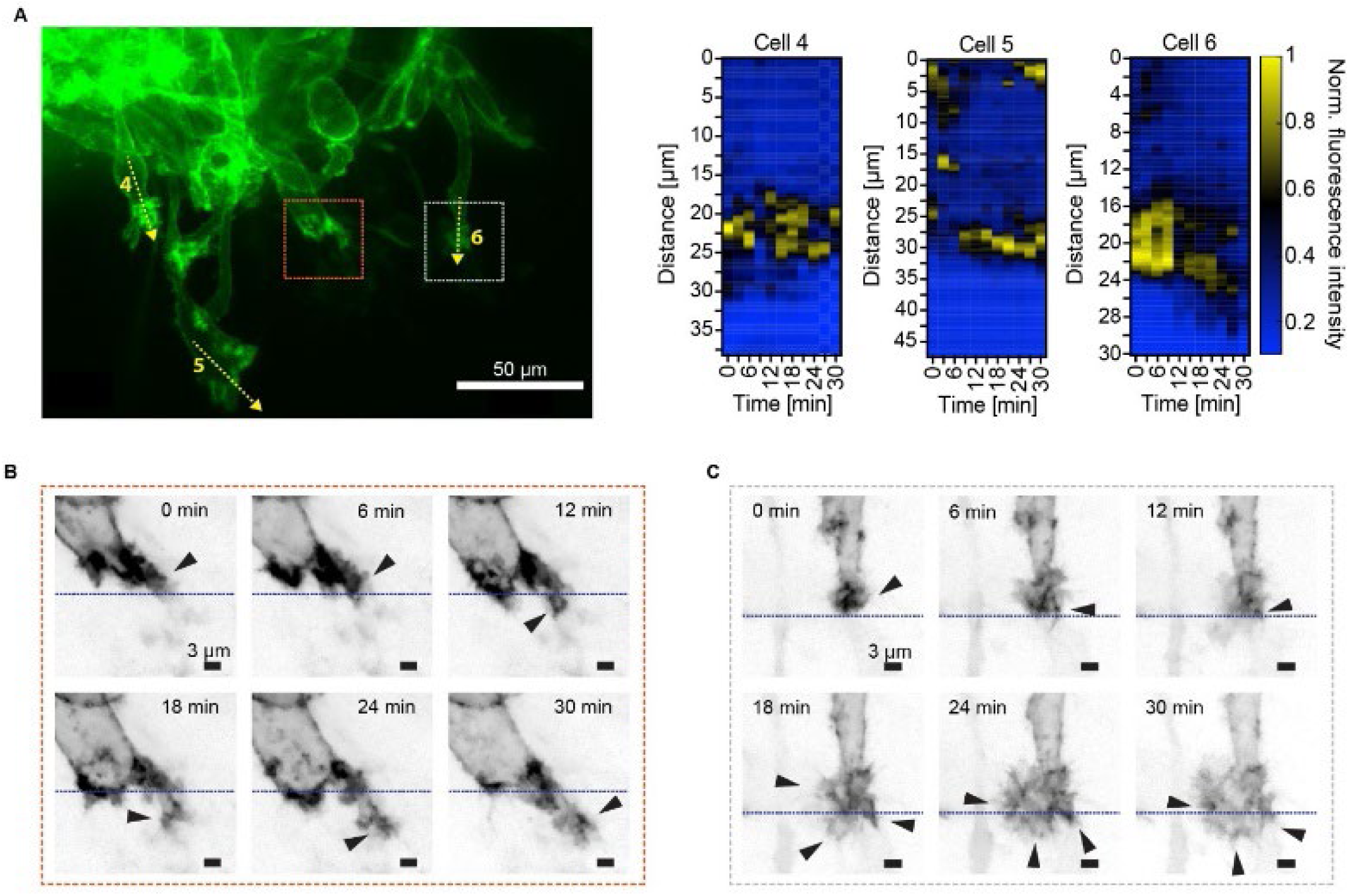
**(A)** Left: Green channel of the image in Figure 5B, with yellow-dotted arrows indicating additionally quantified areas of leading-edge lamellipodia. Right: Heat map kymographs show relative gray fluorescence values of the green fluorescence channel over 30 minutes along a 48 µm line across leading-edge lamellipodia in representative invading cells, in addition to the cells shown in Figure 5G. **(B-C)** Inverted grayscale of B7-H3-mNG fluorescence intensity of two cells from **(A)**, indicated by the dotted bounding box. The images show lamellipodia dynamics over 30 minutes of acquisition, with arrowheads indicating accumulation at the leading edge. The initial position of the leading edge is visualized with the blue dotted line.

**Extended Data 5:**
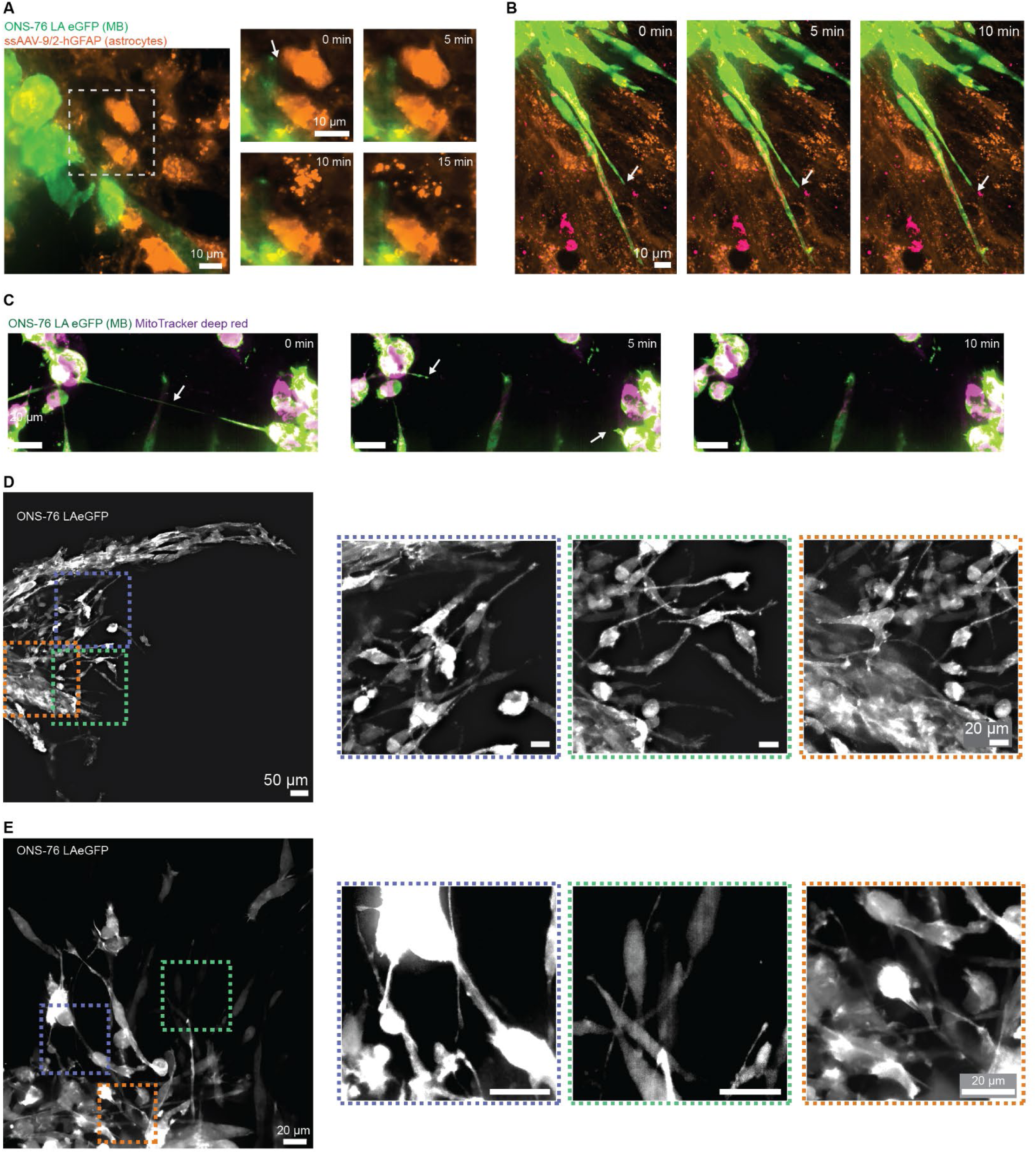
**(A)** Example of an observed disrupting tumor-tumor cell connection. Green: LA-eGFP; Magenta: MitoTracker deep red. **(B)** Still frames of a microtube extending from an ONS-76-LA-eGFP cell towards the TME. The arrows indicate the tip of the microtube. Green: LA-eGFP; orange: GFAP-driven mCherry; magenta: MitoTracker deep red. **(C)** Still frames of an event where tumor-cell contact likely elicited astrocyte death. **(D-E)** Representative images showing live observations of tumor cells interconnected through long, actin-rich protrusions in two additional independent experiments. Images show maximum intensity projections of still images. All scale bars in the close-ups are 20 µm.

