## Supplementary information for "PHIROS: Integrated microfluidic platform for multi-day high-resolution imaging of organotypic slices"

### Methods

#### Platform fabrication

All parts of the platform were fabricated using standard rapid-prototyping techniques. Designs were created in Fusion 360 (Autodesk Inc., San Francisco, CA, United States). An exploded view of the chip and details of the assembly are shown in **Supplementary Figure 1**.

The culturing part of the platform, further referred to as "culture chip", featured an outer diameter of 22.5 mm and was fabricated through micro-CNC milling (Roland SRM-20) of 800 µm polymethylmethacrylate (PMMA) sheets as described elsewhere[1]. The channel width was 700 µm at both the inlet and outlet. To ensure a complete filling of the triangular shaped, 12-mm-wide and 800-µm-deep culture chamber, we incorporated a phase guide of 300 µm height at the outlet channel. The microfluidic channel had a depth of 500 µm. Inlet and outlet diameter were 800 µm. The double-sided adhesive tape (467-7952 MPL, 3M, Rüschlikon, Switzerland) was laser-cut using a $CO_2$ laser (Universal Laser System, Vienna, Austria). After bonding the adhesive tape to the PMMA part, chips were cleaned using a 40% ethanol solution, rinsed with deionized water, and then blow-dried using an air gun. We treated the phase guide with trichloro(1H,1H,2H,2H-perfluorooctyl)silane (448931-10G, Sigma-Aldrich, Buchs, Switzerland), diluted 1:100 in HFE (Novec 7000™, 3M, Rüschlikon, Switzerland), to increase its hydrophobicity. Subsequently, we covered the phase guides with 20 µl of PDMS, prepared in a 1:10 ratio (Sylgard 184, Dow Chemical Company), to protect the phase guide during static culturing in the incubator. Next, the 0.4-µm-pore-size, hydrophilic PTFE biopore membrane (BGCM00010, Sigma-Aldrich, Buchs, Switzerland) was attached to the culture chip via the prepatterned double-sided adhesive. Before tissue culturing, chips were UV-sterilized in a cell culture hood for 20 min each side.

The PMMA and polystyrene parts (125 µm thickness, ST31-FM-000191, Goodfellow Cambridge Ltd., Huntingdon, England) used for connecting and sealing the culture chip were laser-cut using a $CO_2$ laser. The PMMA parts were then annealed for 1h at 80°C and subsequently cleaned by sonication in soap water for 3 min, followed by rinsing with 40% EtOH and DI water. All parts were finally sterilized with UV light for 20 min.

#### Perfusion setup

To perfuse the system, the culture chip was sealed on both sides to create a closed microfluidic network. First, a support component for the culture chip was assembled. The support part was formed by laser-cutting a PMMA sheet of the same thickness as the culture chip and sealed by a 125-µm-thick polystyrene foil. The PMMA and the polystyrene foil were bonded using double-sided tape, which had been previously patterned using a laser cutter (**Supplementary Figure 1B**, step 1). The PDMS of the phase guide in the culture chip was then removed, and the culture chip was inserted into the support part, which yielded an assembly with standard microscopy-slide dimensions and facilitated fluidic connections and handling (**Supplementary**

**Figure 1B**, step 2). Finally, the back part of the culturing chip was sealed with a single-sided adhesive layer consisting of the same double-sided tape and the polystyrene foil, in which openings for fluidic inlet and outlet had been laser-cut (**Supplementary Figure 1B**, step 3). The sealed and assembled chip was then inserted into a custom-made aluminum holder, and a support PMMA structure was used to connect and hold the tubing (**Supplementary Figure 1C**, step 4-6). The sealing procedure was carried out in a sterile cell-culture hood.

**Experimental setup**

Three chips were connected and imaged simultaneously; each chip was connected to its own medium reservoir, which enabled us to use each chip for specific experimental conditions during one imaging assay. The sealed chips were connected to the fluidic system at the Nikon Eclipse Ti2-E microscope (Nikon Europe B.V., Amsterdam, Netherlands) equipped with an X-Light V3 Spinning Disk (CrestOptics S.p.A) module. An environmental stage-top incubator (Life Imaging services, Basel, Switzerland) ensured that the system was kept at 37°C during imaging. A peristaltic pump (ISM931C, Ismatec) was used to drive the fluids. A pulsated flow (7-minute flow intervals at 10 µl/min, 61 minutes pause) was applied.

To avoid uncontrolled gas exchange in the tubings, gas-impermeable tubings were used to connect the medium reservoirs to the oxygenator and the oxygenator to the chip. A combination of different tubings was used to connect the different components: Fluorine rubber material (FPM) tubing (0.5mm ID, 302631, Reichelt Chemietechnik, Heidelberg, Germany), and polyvinyl chloride (PVC) peristaltic pump tubing (0.25 mm ID, N0773117, PerkinElmer, USA) from the medium reservoir to the oxygenator, polyether ether ketone (PEEK) tubing (0.5 mm ID, CIL-1569, IDEX, Northbrook, Illinois) from the oxygenator to the chip. We glued FPM O-rings (1 mm ID, 860404.0003, Brütsch & Rüegger, Urdorf, Switzerland), to the ferrule (P-248X, IDEX) to provide stable and leak-free connections to the chip. For the outlet tubing, we used poly(tetrafluoroethene) or PTFE tubing (0.4 mm ID, S1810-06, Bola, Grünsfeld, Germany). All tubes were flushed with DI water, followed by a 70% ethanol and DI water rinse before use.

The oxygenator consisted of a 50 ml Falcon tube (227261, Greiner Bio-One, St. Gallen, Switzerland) with gas-permeable, 0.4-mm-ID PTFE tubing (S1810-06, Bola, Grünsfeld, Germany) coiled within the Falcon tube. The Falcon tube was partly filled with DI water, which was constantly bubbled with Carbogen to saturate the Falcon environment with humidified Carbogen gas and to provide effective gas exchange through the gas-permeable tube. The tube volume in the Falcon tube was 31 µl.

A custom-made Matlab script was used to control the peristaltic pump and automate the stop-flow perfusion scheme.

**Oxygen Measurements**

The dissolved oxygen concentration in the medium was measured with a fiber-based oxygen sensor (OXR50-OI, Pyroscience, Aachen, Germany). All oxygen characterization measurements were conducted at a flow rate of 10 µl/min and after an equilibration period of 5 min. Oxygen concentrations were measured at the end of the poly(etheretherketone) or PEEK tubing. Measurements were taken for 1 min at a sampling frequency of 1 Hz and repeated thrice per tubing length.

**Cerebellar slice culture**

Animal experimentation for mouse brain tissue harvesting was approved by the veterinary offices of the cantons of Basel and Zurich according to Swiss federal laws on animal welfare and carried out in accordance with the approved guidelines.

Sagittal cerebellar brain slices were prepared as described previously[2]. Briefly, wild-type C57BL/6JRj mice pups were sacrificed at P8–P12 by decapitation following isoflurane anesthesia. Cerebella were harvested and placed in cold Gey's balanced salt solution (G9779, Sigma-Aldrich) containing kynurenic acid (K3375, Sigma-Aldrich), here referred to as GBSSK. Cerebella were then embedded in sagittal orientation in 2% low melting point agarose (A9414, Sigma-Aldrich) prepared in GBSSK. The solidified agarose block was glued onto the vibratome disc (VT 1200 S, Leica) using super-glue. Next, the disc was mounted in the vibratome chamber, filled with cold GBSSK, and 350-µm-thick slices were cut at a speed of 0.2 mm/s and an amplitude of 1.1mm. After removal of excess agarose around the slices, the slices were transferred to petri dishes filled with fresh, cold GBSSK on ice.

The cerebellar slices were then transferred to either standard cell-culture inserts (PICM0RG50, Sigma-Aldrich), or the culture chips, which were placed on cell-culture inserts in 6-well plates with 1 ml of brain slice medium (BSM, described as slice culture medium in[2]). We used fire-polished glass Pasteur pipets to place a maximum of three slices per culture chip or insert. Medium was exchanged daily, except for the day following AAV transduction, when medium exchange was not performed. Slices were cultured in a humidified incubator with 5% $CO_2$ at 37°C.

### AAV transduction

Adeno-associated viruses (AAVs) were obtained from the viral vector facility of the University of Zurich. We used i) ssAAV-9/2-hGFAP-hHBbI/E-GCaMP6f-bGHp(A) at 2 x $10^8$ vg and ii) ssAAV-9/2-hGFAP-mCherry-WPRE-hGHp(A) at 2 x $10^8$ vg to specifically label astrocytes in the cerebellar slices. Viruses were diluted in PBS to achieve the indicated final viral titer concentration in a volume of 5 µl PBS. On DIV4 and DIV6, 5 µl of the prepared virus solution were carefully pipetted on top of each slice during the daily medium exchange. Slices were kept in the cell-culture hood for an additional 10 min following the transduction before bringing them back in the incubator.

### Cell Culture

Medulloblastoma cell lines

ONS-76 LA-eGFP and ONS-76 B7-H3 B7-H3-mNG cells were cultured in RMPI-1640 medium (R0883, Sigma), supplemented with GlutaMax (#35050-038, Gibco), 10% FBS (S0615, Sigma) and 1% penicillin/streptomycin (#15140-122, Gibco). Cells were passaged at 80% confluency and regularly tested for mycoplasma.
Spheroids were formed 48 h before the initiation of the co-culture by seeding 2'500 cells per well in 96 U-bottom well plates (NuclonTM SpheraTM 96-Well, Thermo Fisher Scientific). The spheroids were formed in neurosphere medium, adapted from[3] by using 100 µl medium per well and containing 2% B27 (17504044, Gibco), 1% N2 (17502-048, Thermo Fisher Scientific), 2 µg/ml Heparin (07980, Stemcell Technologies, Basel, Switzerland), 20 ng/ml EGF (PHG0315, Gibco), 10 ng/ml bFGF (PHG0360, Gibco), 1 % penicillin-streptomycin (15140148, Thermo Fisher Scientific) in DMEM-F12 (11330032, Gibco). Cells and spheroids were cultured in a humidified 5% $CO_2$ incubator at 37 °C.

Primary neuronal rat culture

Primary neuronal rat cells were cultured on 24 coverslip-bottom well plates (82426, Ibidi GmbH, Gräfeling, Germany), coated with 0.05% (v/v) poly(ethyleneimine) (day before plating, P3143, Sigma-Aldrich) in borate buffer (Thermo Fisher, check number) and 0.02 mg/ml laminin (30 minutes before cell plating, L2020, Sigma-Aldrich). During laminin incubation, cortices of Wistar rats at embryonic day 18 were dissociated with 0.25% trypsin-EDTA (25200-056, Invitrogen). After 20 min of digestion, the cortices were washed twice with plating medium, then triturated, and the obtained cells were counted using an automated cell counter

(NucleoCounter NC-202, chemometec, Allerod, Denmark). We seeded 75’000 live neural cells per well in 25 µl of plating volume. The plates were incubated at 37°C for 45 min before adding 1 ml of plating medium. The plating medium consisted of Neurobasal (21103-049, Invitrogen) with 10 % horse serum (26050088, Gibco), 0.5 mM GlutaMax (35050-038, Gibco) and 2 % B-27 plus (A3582801, Gibco). After 76 h, 50% of the plating medium was replaced with growth medium consisting of NbActive4 (Transnnetyx, Cordova, TN, USA) with 1% penicillin-streptomycin. 50% of the medium was then exchanged every 3-4 days.

### Generation of fluorescent-protein-expressing cell lines

Origin, maintenance and use of ONS-76 cells in cerebellar slice culture were described in[4]. Stable ONS-76 cells expressing lifeact-enhanced green fluorescent protein (LA-eGFP) or human B7-H3/CD276 fused to mNeon-Green (B7-H3-mNG) were generated by lentiviral transduction. The lentivirus was generated using HEK293 T cells transfected with the lentiviral packaging vectors pPax2 (4.5 µg), the coat protein vector pVSVG (3 µg), and either a lentiviral vector encoding lifeact[41] fused to EGFP or B7-H3-mNG (https://en.vectorbuilder.com/vector/VB220316-1594xpv.html) (7.5 µg). Virus supernatant was harvested 30 h after the transfection and filtered through a 45-µm filter. ONS-76 were transduced in standard 6-well plates with 1 ml of the supernatant supplemented with Polybrene (ThermoFisher Sientific, 1:1000). The transduced cells were cultured in complete growth medium and expanded. Two days after transduction, puromycin selection was started. Selected cells were FACS-sorted to generate a pure population of LA-EGFP or B7-H3-mNG expressing cells.

### Validation of B7-H3-mNG surface expression by flow cytometry

6 x $10^5$ ONS-76 MB cells were seeded in standard 6-well plates and incubated at 37 °C, 5% $CO_2$ for 24 h. Cells were washed in PBS and detached by trypsin incubation. After detachment, trypsin was immediately quenched by the addition of culture medium containing 10% FBS. Cells were fixed in 4% PFA in PBS for 20 min at room temperature and then stained with PE-conjugated anti-human CD276 (1:200, Biolegend #331606) or with the corresponding PE-conjugated isotype control antibody (1:200, Biolegend #400112) for 20 min on ice. Labelled cells were resuspended in 2% FBS in PBS, and signals were acquired using a BD LSRFortessa flow cytometer. Cell fluorescence was analyzed using FlowJO software.

### Co-culture models

Before initiating the co-cultures between DIVs 14-18, the spheroids were checked under the microscope and only round, compact spheroids were selected for implantation. Spheroids were picked up from the 96-well plate with a 10-µl pipette and carefully implanted by placing them on the cerebellar slice or in the center of the well on top of the dissociated neural rat cells. We used a hand-held digital microscope featuring blue excitation LEDs (AM4115T-GFBW, Dinolite) to check if the spheroid had been implanted on the cerebellar slice. The co-culture was initiated 72 h before the start of the perfusion culture and of the imaging assay.

### Live-cell imaging

All live-cell imaging assays were performed on a Nikon Ti-2 equipped with a spinning disc X-Light confocal unit as described in detail in the experimental setup section.

#### <u>Tissue health</u>

Tissue health under perfusion was assessed using live-cell cleaved Caspase-3/7 (Nucview 530, Biotium, Freemont CA, US) and NucBlue (R37605, Thermo Fisher Scientific) to counterstain the nuclei of the tissue slices. We perfused the slices after the start of the perfusion culture, 24 h, and 48 h into the perfusion culture with a staining solution containing 2 drops/mL NucBlue and 2 µM Caspase-3/7 of BSM. Tissue slices were incubated with the staining solution for 1h, before images were acquired at 20X. To determine the imaging settings for the caspase-dye acquisition, we induced apoptosis in tissue slices cultured on a control-chip through incubation with 2 µM Staurosporine for 24 h. The positive control was subsequently stained with the same staining solution containing 2 drops/mL NucBlue and 2 µM Caspase-3/7 and imaging settings were determined. As a real-time, positive control during perfusion culturing, apoptosis was induced by exposing one chip to 40 µM Doxorubicin in BSM and incubation for 2h before imaging. The number of apoptotic cells and the total number of cells per image were estimated using the cell density counting workflow of Ilastik 1.4.0.[5]. For each tissue slice and time point, 3 different areas were imaged. Tissue viability was then calculated by the ratio of caspase-positive cells to the total number of cells.

Calcium imaging

On-chip calcium imaging of the transduced slices was conducted in artificial cerebrospinal fluid (aCSF) and under continuous perfusion to avoid potential clogging of the tubing due to the high salt concentration. The composition of the used aCSF is listed in **Table 1**. All reagents for the aCSF were purchased from Sigma-Aldrich. After image acquisition, the medium was changed back to BSM, and the stop-and-go protocol was resumed. We acquired calcium dynamics in a single plane for 5 min using a 20X (NA 0.45) dry or a 40X (NA 0.8) water immersion objective with an exposure time of 800 ms and 10% resp. 8% laser power.

On imaging day 3, we then exposed the slices with 100 uM carbenoxolone (C4790, Sigma-Aldrich) in aCSF for 1h using the perfusion system. Subsequently, the previously selected image ROIs were reimaged using the same imaging settings.

Medulloblastoma co-culture model

We used 10X (NA 0.45) dry, 20X dry (NA 0.45), 40X (NA 0.8) and 60X (NA 1.0) water-immersion objectives to acquire images of tumor cell invasion and tumor-host interactions. Time lapses at 40X and 60X at high temporal resolution (every 3 resp every 5 min) were acquired during perfusion stops. Overnight, we acquired images at 10X synchronized to the perfusion stops; the images were taken 17 min and 51 min after the flow had stopped.

We exposed both co-culture models to 1 uM cytochalasin-D (CytoD, C2618, Sigma-Aldrich) on the second day of imaging. For the slice co-culture, we used the perfusion system to replace the medium in the culture chamber with medium containing CytoD and started acquiring acute effects on the medulloblastoma cells 10 min after full medium exchange. MB cells co-cultured with primary neural rat cells were exposed to 1 uM CytoD by fully replacing the medium in the wells.

Mitochondria were labelled using the MitoTracker DeepRed™ (M22426, Thermo Fisher Scientific) stain at 1:500 dilution prepared in PBS. The staining solution was carefully placed as a 5 µl drop on top of the slices 1h prior to chip sealing.

**Immunocytochemistry**

Tissues were fixed for 1h in 4% PFA (28908, Thermo Fisher Scientific), diluted in 1X PBS, either i) on a shaker, for slices that were cultured under static conditions, or ii) under continuous perfusion, for on-chip samples. After fixation, samples were cut out from the insert or the chip and stored in PBS at 4°C until staining. Tissues were permeabilized for 5 min in 0.25% Trypsin-EDTA (25200056, Thermo Fisher Scientific), followed by a 1h blocking step in 1X PBS with 3% FBS, 3% BSA (diluted from A1595, Thermo Fisher Scientific) and 0.3% Triton-X100 (X100, Sigma-Aldrich). Primary antibodies were diluted in PBS with 3% FBS, 3% BSA, and samples were incubated overnight under agitation at 4°C. Samples were then washed thrice for 10 min using PBS with 5% BSA. Secondary antibodies were diluted with DAPI (75004, Stemcell Technologies, Basel, Switzerland) in PBS containing 3% FBS and 3% BSA and incubated at room temperature for 3 h. Next, samples were washed three times 10 min with 1X PBS before mounting on 75 mm x 25 mm glass slides using ProLong Glass antifade mounting (P36984, Thermo Fisher Scientific) medium and covering with a #1.5 coverslip. Mounted samples were imaged with a Nikon Ti2 spinning-disc confocal using a 10X dry and 40X water-immersion objective.

The following primary antibodies were used: rabbit anti-Calbindin (1:1000 dilution, ab108404, abcam), guinea-pig anti-GFAP (1:2000 dilution, 173 308, Synaptic Systems), chicken anti-NeuN (1:500 dilution, ABN91MI, Fisher Scientific), rabbit anti-HIF-1α (1:500 dilution, ab179483, abcam) and rabbit anti-S100β (1:100, ab52642, abcam). As secondary antibodies we used Alexa Fluor donkey anti-rabbit 647 (A21206, Thermo Fisher Scientific), Alexa Fluor donkey anti-chicken 488 (A78952, Thermo Fisher Scientific), and Alexa Fluor goat anti-guinea pig 555 (A21435, Thermo Fisher Scientific) at 1:200 dilution. Hoechst was added to the secondary antibody solution at a dilution of 1:200.

**Image Analysis**

Tissue integrity

Purkinje cell somas were segmented in stacks of 10 µm, acquired at 0.4µm z-step size using Imaris v10.0.1 to determine the volume of each Purkinje cell soma and the number of cells per ROI. To normalize the number of cells with respect to the area of the Purkinje cell layer (PCL), we created maximum intensity projections of the ROIs. Next, boundary boxes of the PCL were drawn in ImageJ (2.14/1.54f) using the polygon tool and measured, and the number of segmented cells was divided by the measured PCL area.

Hypoxia

Positive controls to determine correct imaging settings of hypoxic cerebellar slices were generated by submerging slices cultured on inserts in either 1 mL or 2 mL of medium, mimicking conditions during upright imaging. We used Imaris v10.0.1 to assess hypoxic cells in a 10-µm stack acquired in the middle of the tissue slice. Nuclei were segmented as surfaces, while HIF-1alpha expression was segmented using the spots tool. To determine the number of truly hypoxic cells, the number of HIF-1alpha spots within nuclei surfaces was calculated. The increase of hypoxic cells under the different conditions was then calculated as an x-fold increase in hypoxic cells with respect to the average percentage of hypoxic cells detected in the standard ALI culture on tissue culture inserts. In total, we cultured 8 slices from 2 independent experiments under pulsatile flow and analyzed images from 2-3 ROIs per slice.

Calcium imaging

We used AQuA2 to analyze the calcium dynamics in transduced astrocytes[43]. Preprocessing of the acquired files included the removal of the first 21 frames of each acquisition, since we observed a strong intensity decay within the whole ROI, likely due to activation of Calcium channels by the laser light[6]. Batch processing of all acquired files was performed using Matlab (R2024a, The Mathworks, Inc., US) after confirming the settings using three to four different acquisitions. The following parameters were used for 40X acquisitions: median filter 1.5, Gaussian filter radius 0.5, intensity threshold 2.8, minimum duration 4, minimum size (px) 15, minimum seed size 0.01, Z score 3.5, maximum dissimilarity 0.7, minimum source size 0.01, sensitivity level 7. For 20X acquisitions the parameters changed to the following: median filter 3, Gaussian filter radius 0.7, intensity threshold 2, minimum duration 5, minimum size (px) 18, minimum seed size 0.01, z score 1.8, maximum dissimilarity 0.7, minimum source size 0.01, sensitivity level 8. After the batch processing, results of detected events were further filtered to include only events with an event size between 10-2500 $\mu m^2$ for 40X images and 20-2500 $\mu m^2$ for 20X images as well as an event duration between 1.6 and 60 s. To compare the effect of carboxenole (CBX) treatment on velate astrocytes, we further filtered the AQuA detected events and included events larger than 150 $\mu m^2$.

Cell tracking and particle image velocimetry:

To estimate cell velocity and displacements of tumoral cells and astrocytes, we analyzed the overnight timelapse images using the PIVLab[7] Matlab app for particle image velocimetry. The acquired series of time-resolved images were pre-processed by subtracting the mean intensity across the image series and then processed using a multi-pass FFT window deformation method to extract the velocity fields. A four-pass cross-correlation approach was applied, starting with an interrogation area of 312 pixels and a step size of 100 pixels. In each subsequent pass, both the interrogation area and step size were reduced by 50%. The mean and standard deviation of velocity across the 12-hour acquisition period were visualized as a heatmap.

Cell morphological analysis

60X-magnification, z-stack timelapse acquisitions were first preprocessed by denoising, followed by maximum intensity projection and local-contrast enhancement in NIS Software. These pre-processed timeline acquisitions were subsequently analyzed in Fiji using the plugin Trackmate-Cellpose[8] using the pre-trained model Cyto3. Morphological parameters were extracted for cells whose segmentation and tracking could be carried out for 15-60 continuous minutes. Cell-area variability was calculated as the ratio of the standard deviation of the area during the observation period over the mean area.

Filopodia quantification

60X-magnification acquisitions were first preprocessed as described before for the morphological analysis. ROIs containing single cells or small clusters of cells were then extracted and analyzed in Fiji using the Filoquant plugin[9] to calculate edge and filopodia lengths.

Quantification of astrocyte morphology

To assess astrocyte morphological dynamics across the three-day imaging period, we employed an adapted version of the previously established semi-automated MicrogliaMorphologypipeline[10] implemented in FIJI. The pipeline is designed to process single-channel 3D image stacks or individual two-dimensional images containing clearly delineated somata and processes, permitting its application to glial populations beyond microglia. The macro-based framework integrates functionalities from the BioVoxxel Toolbox and FracLac plugins, which are not fully accessible through the native ImageJ macro language. As a result, plugin-dependent steps were executed manually following macro-generated prompts, while all remaining image preprocessing, feature extraction, and quantitative analyses were automated. User input was restricted to specifying input and output directories and, where appropriate, enabling batch processing.

Single-channel images acquired at 20X magnification were exported. Astrocyte segmentation was performed using the Huang2 thresholding algorithm with defined lower and upper thresholds of 252 and 1133, respectively. The thresholds were defined based on particle selection, excluding image artifacts. Binary masks were subsequently subjected to FracLac analysis, and foreground pixel counts were converted to cell area for quantitative morphological evaluation. In total, four independent chips were analyzed.

Tumor microtubes and mitochondria dynamics in the slice-culture model

Long, actin-rich protrusions were manually measured in Imaris (v 10.2.0) using the filament tracker tool. The thickness of the tumor microtubes were adjusted and measured based on the average thickness in the 3D visualization. Start and endpoint for the length of tumor-tumor connections was determined as the point where the cell soma ended and the protrusions started. To measure the length of connections between tumor cells and the microenvironment, the tip of the protrusion was determined as the endpoint and the start point was determined analogously to the tumor-tumor connections. For the final analysis, we only included connections with a thickness between 0.5- 2.5 um and a length of minimum 10 um based on definitions found in literature[11].

Mitochondria within the microtubes were tracked manually using the spot tracking tool in Imaris (v 10.2.0) by defining the centre of mass of the single mitochondria frame by frame.

<u>Quantification of B7-H3-mNG localization:</u>

A 10-µm-long line with a width of 100 pixels was drawn across the leading-edge area in the direction of migration or perpendicularly across cell-cell contacts in the B7-H3-mNG fluorescent channel in ImageJ. The lines were centered to the leading edge for lamellipodial B7-H3 or to the cell-cell contacts. The profile of the line was plotted using ImageJ, and the grey values were determined every 0.09 µm. The sum of all grey values was calculated, and the percentage of total grey values calculated every 0.09 µm was plotted. An unpaired t-test with individual variance for each row was performed. The false discovery rate was calculated using a two-stage step-up approach[12].

**Data Visualization and Statistical Analysis**

Calcium imaging events were visualized using a custom Matlab script to filter events and visualize events according to occurring frequency distributions.

For the remaining figures, GraphPad Prism (v 10.5.0) was used to visualize data and perform statistical analysis. The data are presented as mean +/- s.e.m.. Distributions were first tested for normality and lognormality, followed by either an unpaired, one-sided t-test or an appropriate one-way ANOVA based on data distribution. Statistical significance was defined as p-values < 0.05.

**Table 1**: Composition of the used aCSF for Calcium imaging

| **Reagent** | **Working concentration** |
|---|---|
| **NaCl** | 126 mM |
| **KCl** | 3.5 mM |
| **$NaH_2PO_4.H_2O$** | 1.2 mM |
| **$MgCl_2.6H_2O$** | 1.3 mM |
| **$CaCl_2.2H_2O$** | 2 mM |
| **D-Glucose** | 11 mM |
| **$NaHCO_3$** | 25 mM |

### Supplementary Figures

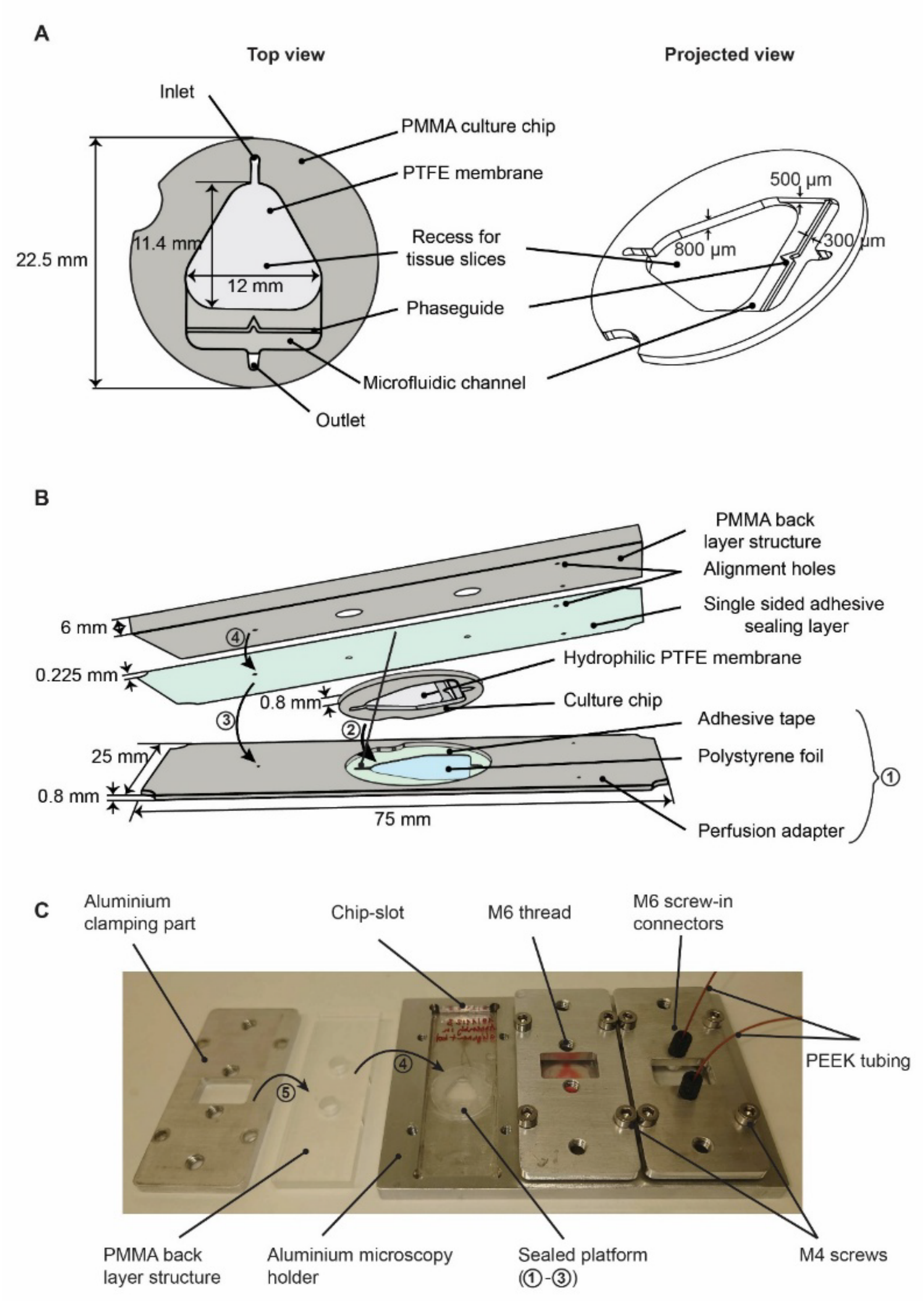


**Supplementary Figure 1: (A)** Top view and projected view of the culture chip with channel dimensions. **(B)** Exploded view of the sealed platform. The perfusion adapter (1) is assembled before initiating the sealing procedure. The culture chip is then snap-fitted into the perfusion adapter (2) and sealed from the back (3-4). **(C)** Photograph of the pieces used to clamp the sealed platform in the microscopy holder and the fluidic connections.

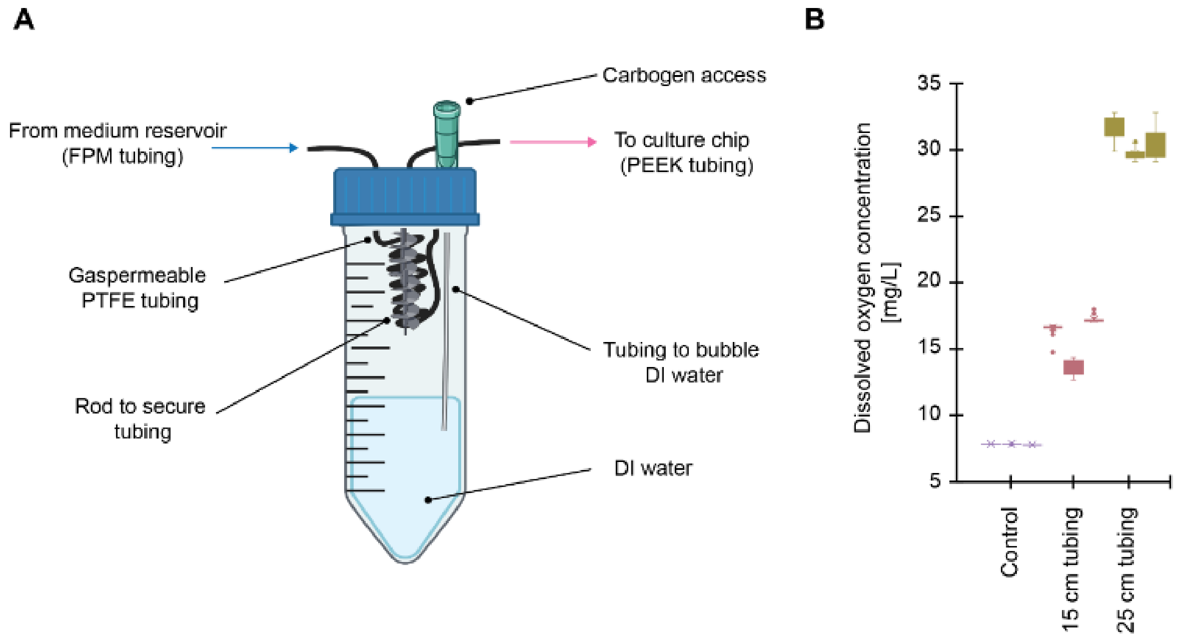


**Supplementary Figure 2**: **(A)** Details of the oxygenator used to buffer and oxygenate the culture medium before entering the chip. A gas permeable tygon tubing was coiled in a 50-ml falcon tube, and the water in the falcon tube was bubbled with a 95% $O_2$, 5% $CO_2$ mixture. **(B)** Measured dissolved oxygen concentration in water at a flowrate of 10 µl/min with tubing lengths of 15 cm or 25 cm.

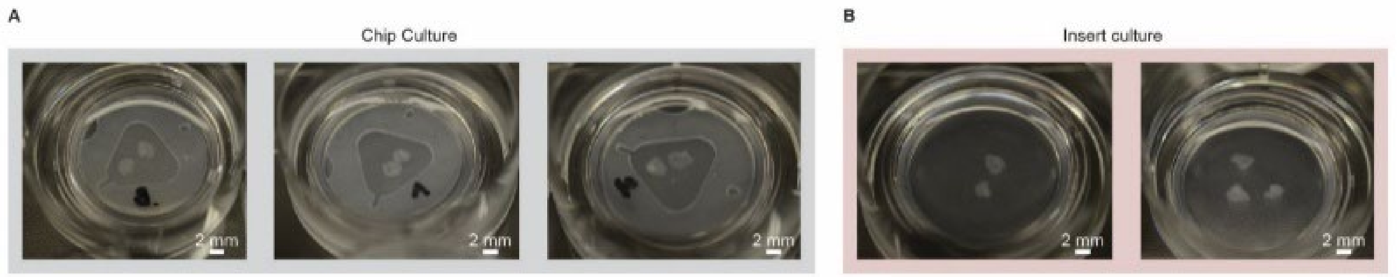


**Supplementary Figure 3:** Transparency of cerebellar slices cultured in dual membrane cultures on chip (**A**) or on standard tissue culture inserts (**B**).

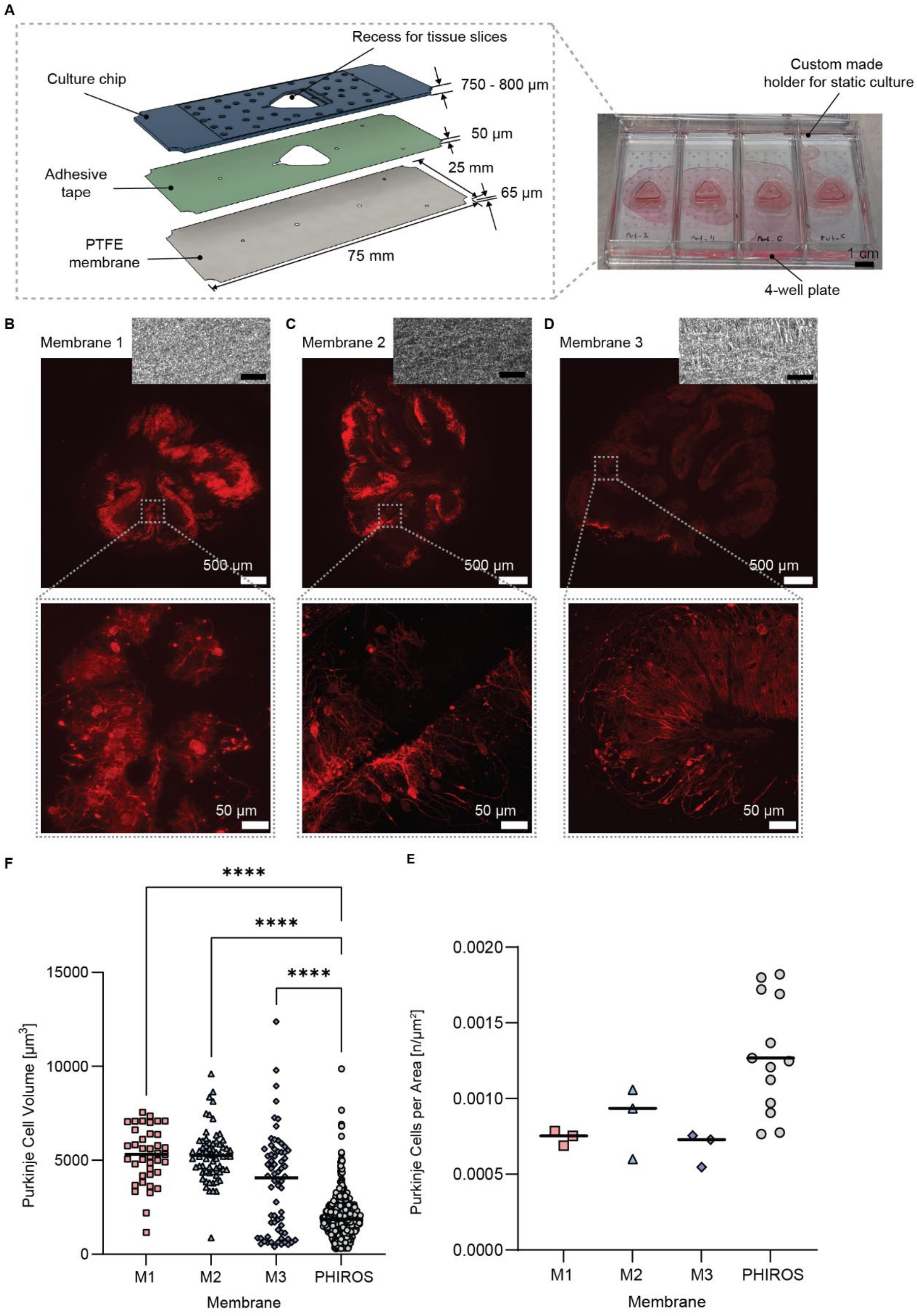


**Supplementary Figure 4**: **(A)**: Schematic of a previous version of the culture chip design with glass slide dimensions (25 mm x 75 mm) using the same adhesive tape and hydrophilic 0.4 µm PTFE membranes from various manufacturers. This previous prototype was placed on a custom-made holder in a 4-well plate during static culturing (photo on the right). **(B-D)** 10X images and 40X zoomed-in images of Purkinje cell layer reorganization on previous platform prototypes fabricated with different hydrophilic PTFE membranes; insets at the top show the structure of the corresponding membrane with 150X magnification, scale bars 10 µm. The Purkinje cell layer was disrupted and showed larger Purkinje cell somas **(E)** and decreased Purkinje cell density **(F).**

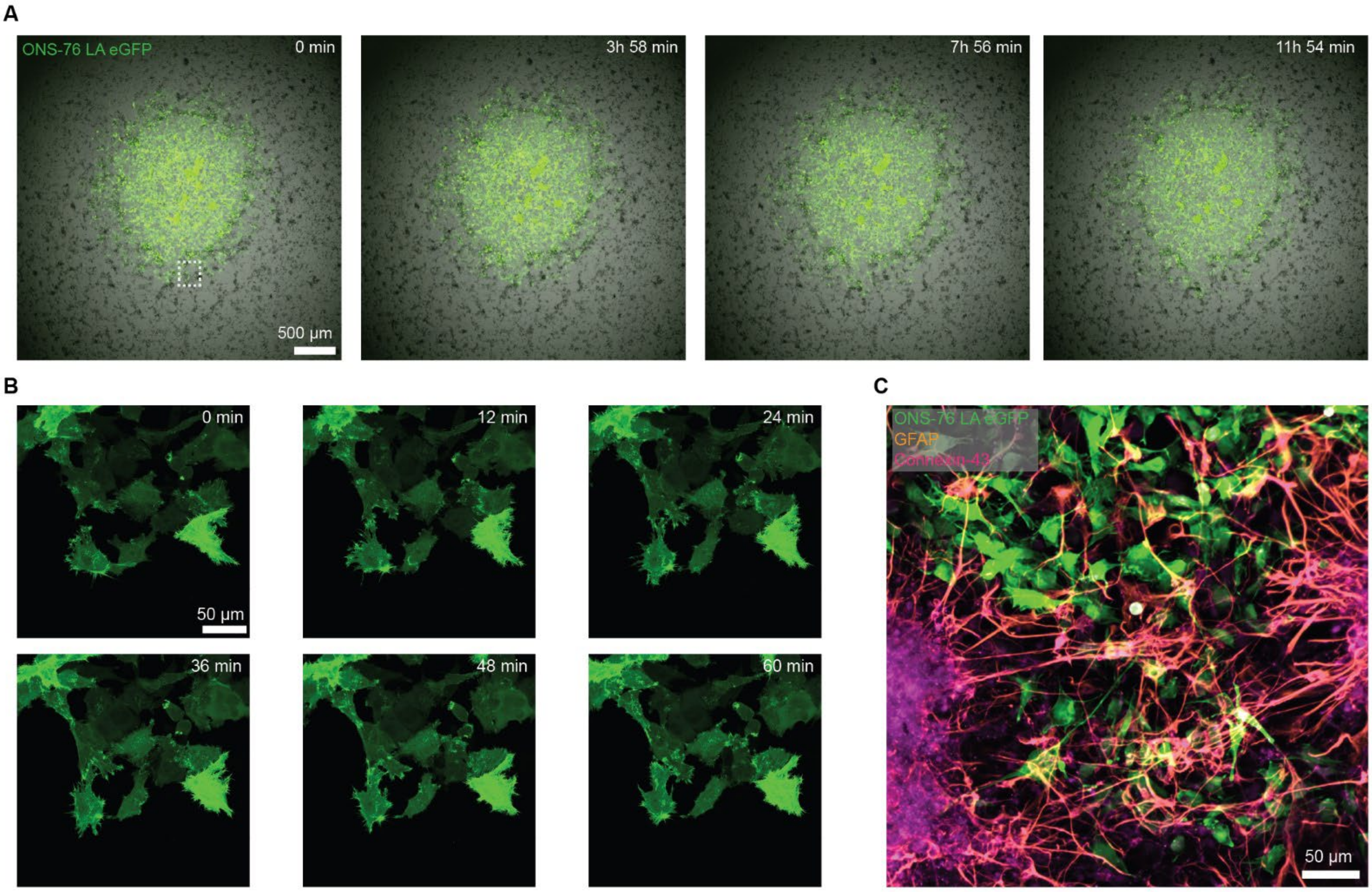


**Supplementary Figure 5:** ONS-76 LA eGFP cells moving on a layer of dissociated neural cells. **(A)**: Progression of the spherical movements during an overnight image acquisition at 4X magnification. **(B)** Tumor cell motility of the inset in B as acquired at higher temporal and spatial resolution. **(C)** Representative maximum intensity projection of a immunofluorescence-stained co-culture of tumor cells and neural dissociated cells. Green: life- actin cytoskeleton of the tumor cells, orange: astrocytes stained by anti-GFAP, magenta: gap junctions of other neural cells stained with anti-connexin-43.

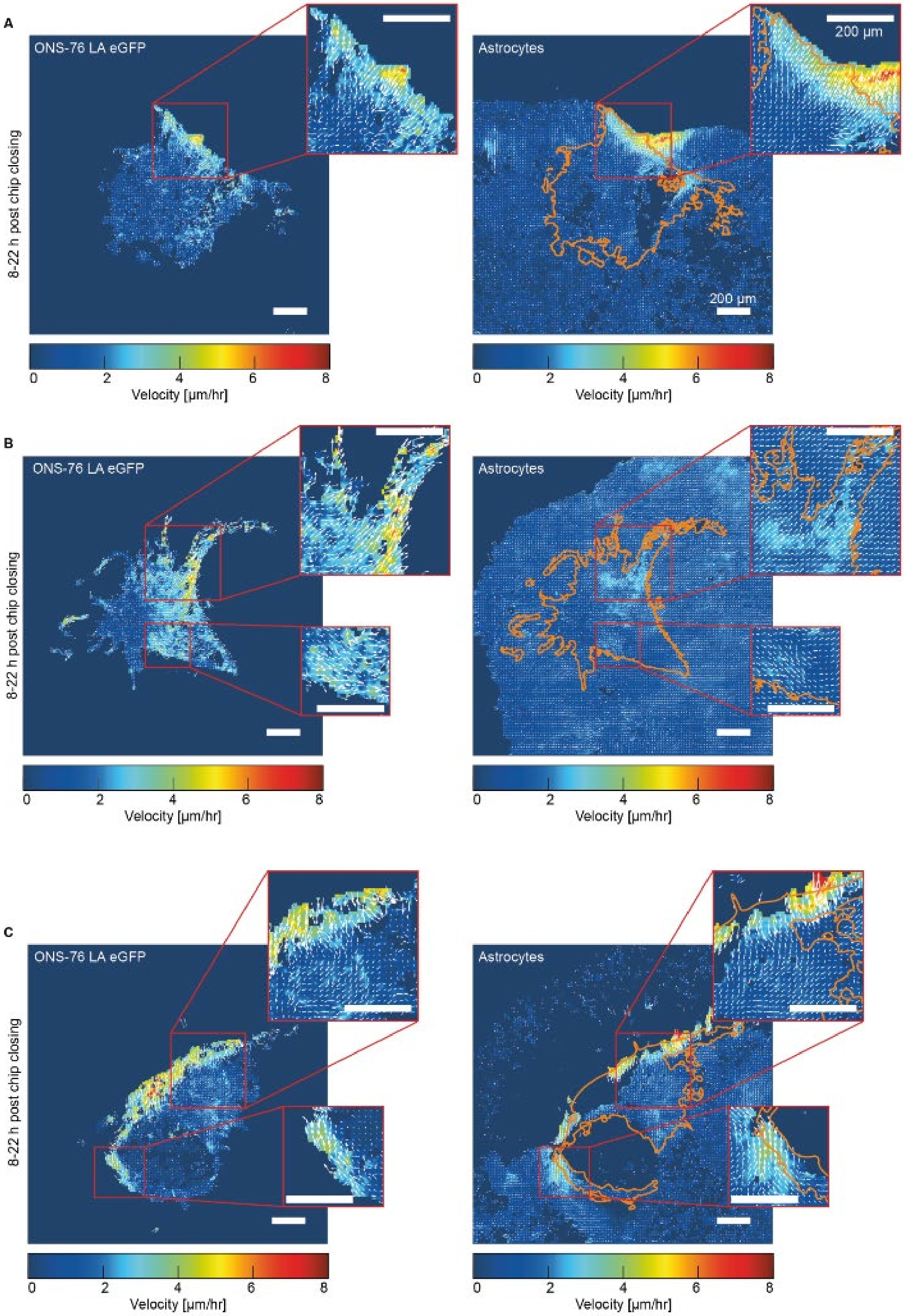


**Supplementary Figure 6**: Mean velocities calculated using PIVLab for both LA-eGFP (left side) and GFAP-driven mCherry (right side). (**A-C**) show three different, independent experiments. Time refers to hours after platform sealing. Scale bar= 200 µm.

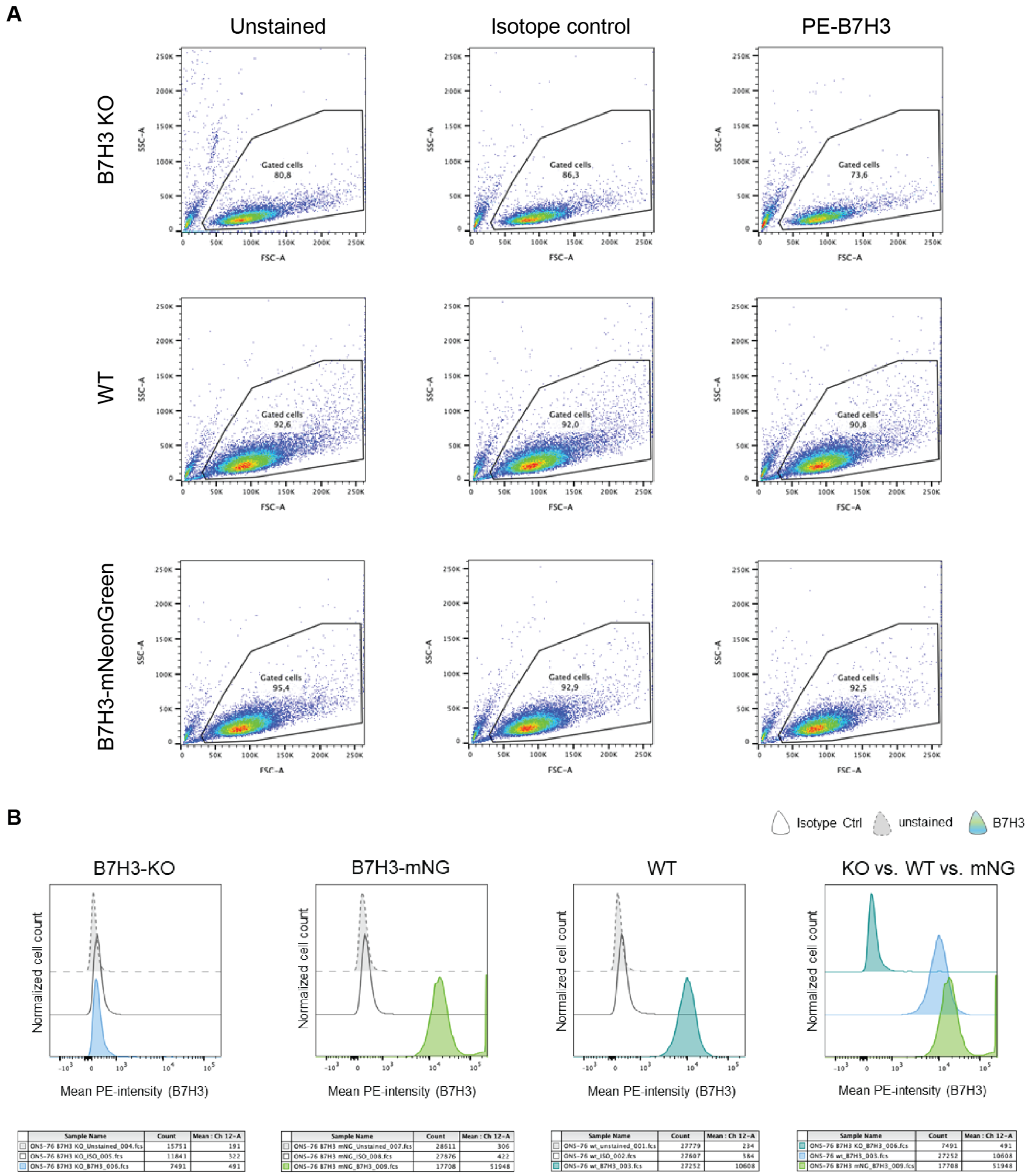


**Supplementary Figure 7**: (**A)** Density plots of flow cytometry analysis of ONS-76 B7H3 KO cells (top row), ONS-76 wildtype cells (middle row) and ONS-76 B7H3 mNeonGreen overexpressing cells (bottom row) of the unstained cell population, the IgG1 mouse isotope control population, and the B7H3 stained population. (**B)** Histograms of the same cell populations confirming the stable expression of B7H3 in the genetically modified ONS-76 population as evidenced by the overlap with WT cells co-stained with B7H3 in the rightmost panel.
